# Interferon-γ promotes SMAP production by cytotoxic T lymphocytes in a thrombospondin-4 dependent manner

**DOI:** 10.64898/2026.06.12.731857

**Authors:** Omnia M. Khamis, Mohammad Shkeir, Claudia Schirra, Szu-Min Tu, Chiara Cassioli, Abed Alrahman Chouaib, Eve Lecomte, Lena Schönecker, Michael L. Dustin, Hsin-Fang Chang, Cosima T. Baldari, Ute Becherer

## Abstract

Cytotoxic T lymphocytes (CTLs) eliminate infected and cancerous cells by exocytosing cytotoxic granules, either as single-core granules (SCGs) releasing diffusible Granzyme B and Perforin, or as multi-core granules (MCGs) releasing these effectors as thrombospondin-1/4-encapsulated supramolecular attack particles (SMAPs). How CTLs differentially deploy these granule types remains unclear. We demonstrate that prolonged *in vitro* expansion and restimulation selectively enhance SMAP release, correlating with increased MCG maturation and thrombospondin-4 expression. Using high-resolution imaging, we identify fusion-competent MCG intermediates lacking SMAPs but releasing granzyme B diffusively. Mechanistically, interferon-γ upregulates thrombospondin-4, driving MCG maturation and SMAP biogenesis, enhancing late-phase CTL killing efficiency against resistant targets. Consistent with this, *THBS1* and *THBS4* transcript levels appear elevated in melanoma-infiltrating CTLs compared with those during acute adenovirus infection. These findings define a stimulus-dependent, interferon-γ–driven pathway tailoring CTL responses to chronic pathology and highlight opportunities for SMAP-targeted immunotherapies.

## Introduction

Cytotoxic T lymphocytes (CTLs) are a key component of the adaptive immune system for fighting against virus infected cells and cancer cells. They do so by specifically recognizing their target cells with their T cell receptor, triggering the formation of an immunological synapse (IS) that polarizes the killing arsenal of the CTL toward the target cell ^1^. CTLs kill the target cells primarily by releasing the content of their cytotoxic granules (CGs) namely Granzymes (Gzm) and Perforin ^2, 3^. This is accompanied by the release of Interferon-γ (IFN-γ) ^4, 5^, and exosomes ^6, 7^ both having cytotoxic activities. Finally, exposure of FAS ligands (FasL) at the IS also contributes to the killing of the target cells ^8, 9, 10^.

In CTLs and natural killer (NK) cells, CGs exist in two distinct morphological classes: single-core granules (SCGs) and multi-core granules (MCGs). SCGs contain one large electron-dense core, which includes GzmB complexed with the proteoglycan serglycin ^11^, and release their content exclusively in a freely diffusible form. In contrast, MCGs harbor multiple smaller electron-dense cores composed of condensed GzmB and perforin, surrounded by a glycoprotein shell containing primarily Thrombospondin-1 and −4 (TSP-1 and TSP-4) ^12, 13, 14^. These stabilized structures correspond to supramolecular attack particles (SMAPs) ^12^. Although the cores of SCGs and SMAPs share compositional similarities, the TSP-rich shell of SMAPs is thought to encapsulate and stabilize a condensed, positively charged GzmB–perforin complex with the negatively charged serglycin ^11^, thereby providing an alternative, non-diffusible mode of effector delivery.

The biogenesis of SMAPs is TSP-dependent, whereby TSP-4 is required at an early stage promoting TSP-1 localization to MCGs ^14^. Both are required for SMAP stability and their killing efficiency ^12, 14^. In addition to the SMAPs, MCGs also contain diffusible Perforin and Gzms, IFN-γ, and exosomes ^4, 6^. MCG and SCG fusion with the plasma membrane strictly depends on the IS formation ^6, 12, 13, 15^ and CTLs appear to preferentially fuse MCG over SCG ^6, 13^. Given their broader and more diverse cargo, MCGs likely support multiple effector modalities, including direct cytotoxicity and paracrine signaling, whereas SCGs may primarily mediate diffusible cytotoxic responses. This points to functional plasticity in CTLs and NK cells, whereby differential deployment of MCGs and SCGs could tailor effector outcomes to specific contexts. However, it remains unclear whether the relative abundance of these granule populations is stimulus-dependent or whether their secretion can be selectively regulated.

Which type of CGs is exocytosed by CTLs might depend on the stimulus strength at the IS or could be regulated by cytokines. CTLs are exposed to a large array of cytokines during target cell elimination ^16, 17^. They express receptors for a wide variety of interleukins (ILs), IFN-α, β and γ, tumor necrosis factor-α (TNF-α), and other bioactive molecules. Furthermore, they secrete IL2, TNF-α, IFN-γ, and others depending on the CTL subtype, that have an autocrine and paracrine activity ^16, 18, 19, 20^. While IL-2 and IFN-α, β are primarily involved in CTL expansion and differentiation from naïve to effector cells, other ILs are involved in memory cell formation (IL-2, IL-7) ^17, 21^ or regulate CD8^+^ lymphocyte activation states (IL10, ^22^). TNF-α and IFN-γ are well known to induce cell death in the target cell ^4, 23, 24^ but especially IFN-γ is also known for regulatory roles on the CTLs. It controls CD8^+^ cell expansion ^25, 26^, their differentiation to CTLs ^27^, immunodominance ^28^ and their mobility and infiltration rate in tumor ^5, 29^.

In this study, we investigated whether the secretion of SCGs and MCGs in murine CTLs is differentially regulated. To address this, we combined live–cell total internal reflection fluorescence (TIRF) microscopy to monitor granule exocytosis in real time with ultrastructural analyses by transmission electron microscopy (TEM) and correlative light and electron microscopy (CLEM). Functional killing assays were used to assess how granule maturation impacts CTL effector function. Using these approaches, we show that prolonged culture and restimulation selectively enhanced SMAP release as CTLs exhibit an increasing number SMAP–containing MCGs. We further identified MCG intermediates containing intraluminal vesicles, GzmB but lacking SMAPs, which were exocytosis competent and released GzmB in a diffusive manner. Notably, we identify IFN-γ as a key regulator of this maturation process, acting to upregulate TSP-4 expression and thereby promote SMAP biogenesis. Functionally, CTLs maintained longer in culture displayed enhanced cytotoxicity. By defining the molecular pathways controlling SMAP formation and release, we provide a framework to harness SMAPs for advanced therapeutic applications.

## Results

### Stimulus strength does not determine secretion type in CTLs

Immunological synapse formation on supported lipid bilayers (SLBs) had been shown to depend on the amount of anti-CD3ε monoclonal antibody integrated in the lipids ^30^ and we wondered whether it also impacted the type of CGs that CTLs release. To test this, we seeded wild type (WT) mouse CTLs, which were transfected with GzmB-pHuji about 16 h prior to the experiment, onto SLBs containing 250 ng/ml ICAM-1 and either 5, 10 or 20 µg/ml anti-CD3ε monoclonal antibody. We visualized CG exocytosis by TIRF-microscopy for 15 min at 10 Hz and at 20°C and observed two types of secretion. The CGs suddenly brightened up upon fusion with the plasma membrane due to the opening of the fusion pore, deprotonation of the CG lumen and dequenching of the pH sensitive pHuji (Fig. S1A, B). Then two different scenarios where observed. Either GzmB-pHuji diffused quickly away generating a cloud of fluorescence (Fig. 1A top, Fig. S1B top), which is probably associated with SCG exocytosis ^13^. Otherwise, GzmB-pHuji positive particles, i.e. the SMAPs, remained visible after part of GzmB-pHuji had diffused away (Fig. 1A bottom, Fig. S1B, bottom). This is likely due to MCG exocytosis ^13^. Based on the different GzmB-pHuji diffusion properties, we refer to the former as cloud release and to the latter as SMAP-release. Of note, the time taken for IS formation and the percentage of cells that secrete were unaffected by the strength of the stimulus. This result is inconsistent with previous data ^30^. Furthermore, the stimulus strength had no impact on the type of secretion (Fig. 1SE, F). By comparing the two studies, we found that Estl, Blatt ^30^ only used cells that had been maintained for 8–9 days in vitro after activation, whereas we used cells that were expanded for 5–8 days. This discrepancy led us to hypothesize that the duration of culture might be a critical determinant of secretion mode, prompting us to systematically investigate how time in vitro influences CTL secretion type.

**Figure 1:**
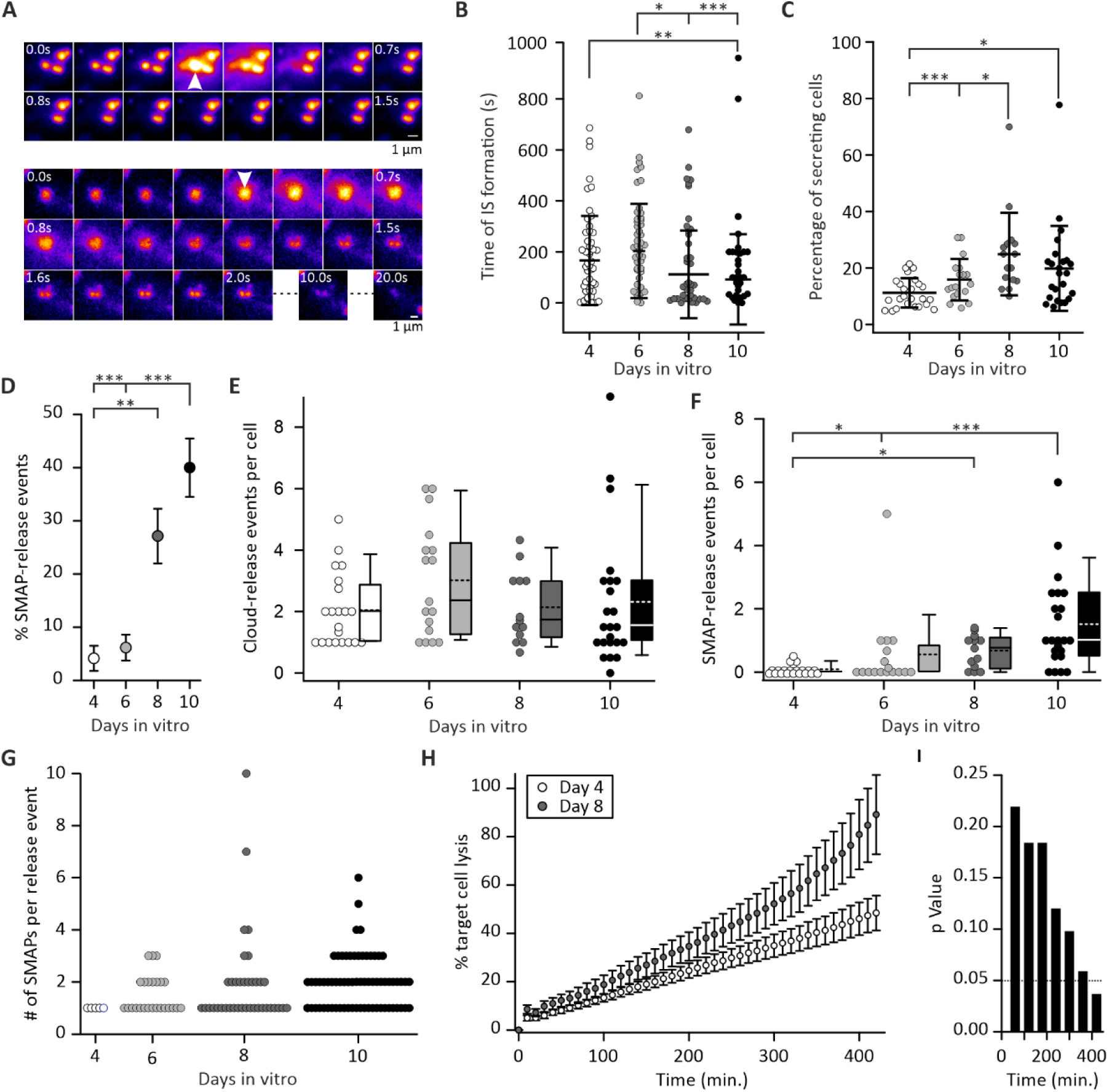
Prolonged CTL culture shifts fusion type from cloud-release towards SMAP-release. **A.** Snapshots of an exemplary cloud- (top) and SMAP-release (bottom) event. The CTL displayed in the top and bottom rows were maintained for 4 and 10 days after activation in culture, respectively, prior to the experiment. Cytotoxic granule (CG) exocytosis was induced by seeding the CTLs overexpressing GzmB-pHuji on activating supported lipid bilayer (SLB) containing 10 µg/ml anti-CD3ε monoclonal antibody. Acquisition was performed at 10 Hz by total internal reflection fluorescence microscopy (TIRFM). Exocytosis is marked by an arrow. Scale bar is 1 µm. **B-C.** Scatter dot plots of (**B**) the time of IS formation and (**C**) the percentage of secreting CTLs at different days during their in vitro expansion. Error bars are SDs. **D.** Diagram representing the mean percentage of SMAP-release per CTL at different days during their in vitro expansion. Error bars are SEMs. **E-F.** Box and scatter dot plot of the normalized number of cloud- (**E**) and SMAP-release (**F**) per CTL and video over their time in culture. Note that, due to space constraints, markers for 0 SMAP-release at day 4 are staked on two rows in ***F***. Stippled line in the box plot represents the mean and the solid line the median. **G.** Scatter dot plot showing the number of SMAPs released per exocytosis event over CTL time in culture. N_mice_ = 6; n_video_= 21, 17, 13 and 23; n_cell_ = 49, 45, 51, and 48 for 4, 6, 8 and 10 days in vitro, respectively. Significance was tested with Kruskal-Wallis one-way ANOVA on rank followed by Dunn’s multiple comparison. * *p*<0.05, ** *p*<0.01, *** *p*<0.001. **H.** Real-time calcein release-based killing assay. Day-4 (white) and 8 (gray) CTLs were co-cultured with B16V cells at an E:T ratio of 10:1, and killing was measured every 10 min for 7 h. Shown is the mean with error bars as SEMs. N_mice_=3, n_repeats_=9. **I.** Graph representing the t-test p value comparing the difference of killing percentage between day-4 and 8.

### Prolonged post-activation expansion of mouse CTL promotes SMAP release

To assess the effect of time after bead activation on their secretion capability more thoroughly, WT CTLs were expanded for 4 to 10 days in vitro prior to the experiment. The cells were then transfected with GzmB-pHuji and about 12 h later they were seeded onto SLBs containing 250 ng/ml ICAM-1 and 10 µg/ml anti-CD3ε monoclonal antibodies to stimulate CG exocytosis. These SLBs were used for the rest of the work and are referred to as activating SLBs. The exocytosis was recorded by TIRF-microscopy. At 4 days post-activation we observed readily cloud-release exocytosis events (Fig. 1A top, Video S1), whereas at 10 days we observed more SMAP-release events (Fig. 1A bottom, Video S2). An in-depth analysis of the TIRF-microscopy videos revealed that the overall behavior of CTLs changed with increasing time in culture. The time required for IS assembly decreased significantly with increasing days post-activation, while the proportion of CTLs that secreted GzmB-pHuji increased significantly. This indicates that CTLs became more efficient over time. More interestingly, the proportion of SMAP-release events over the total release events increased by more than a factor of 10 between day 4 and 10 (Fig. 1D). This increase was not attributable to a decline in the amount of cloud-release, which remained stable throughout all days after activation (Fig. 1E), but rather to a significant increase in the number of SMAP-releases (Fig. 1F). The number of SMAPs released per exocytosis event remained constant at an average of 1.6 SMAPs across all days post-activation (Fig. 1G).

We then assessed if this enhanced SMAP-release was due to a change in CTL subset during cell expansion by subjecting the cells to flow cytometry analysis (Fig. S2). The degree of differentiation of the CTLs was determined using CD44 and CD62L surface markers to define naïve (CD44-/CD62L-), central memory (CD44+/CD62L+), and effector memory (CD44+/CD62L-) CD8⁺ T cells ^31^. We found a steady decrease of the naïve and central memory CD8⁺ T cells subset over time in culture and an increase of the effector memory CD8⁺ T cells subset (Fig. S2B). These phenotypic shifts were not statistically significant and insufficient to explain the massive increase in SMAP secretion, suggesting that time in culture promotes a deeper functional reorganization of the CTL killing arsenal beyond traditional differentiation states.

Finally, to investigate whether increased SMAP release correlates with enhanced killing efficiency, we employed a real-time calcein release-based killing assay (Fig. 1H). Experiments were performed using day-4 and 8 WT CTLs and B16V target cells, with an effector-to-target (E:T) ratio of 10:1. B16V cells were selected over P815 cells based on the hypothesis that SMAPs are particularly critical for eliminating hard-to-kill cancer cells ^32, 33^. At early time points, killing efficiency was comparable between day-4 and 8 CTLs. However, day-8 CTLs exhibited significantly greater killing efficiency at later stages of the assay (Fig. 1H). After 7 hours, day-8 CTLs demonstrated a marked improvement in target cell lysis compared to day 4 CTLs (Fig. 1I). This trend was also observed at an E:T ratio of 1:1 (Fig. S2C, D), although the difference was less pronounced than at 10:1. Given previous findings suggesting that SMAPs are capable of latent target cell killing ^12, 14^, our results indicate that the enhanced killing efficiency of day 8 CTLs is tightly linked to their increased SMAP release leading to improved killing late in encounter with targets.

### Extended CTL expansion increases multi-core granules number and size and enlarges SMAP

Since SMAP-secretion is most likely due to MCG fusion with the plasma membrane, its increase could be explained either by a change in the release machinery in favor of MCG, a promotion of MCG trafficking towards the IS or an enhanced MCG biogenesis over time in culture.

To investigate whether time in culture affects MCG biogenesis, we compared CTLs at 4 and 8 days post-activation at the ultra-structural level using transmission electron microscopy (TEM). These time points were chosen as day-4 CTLs displayed the lowest number of SMAP-release events, whereas at day 8 the number of SMAP-release events was substantially increased while preserving cell viability. CTL were seeded on anti-CD3ε antibody coated sapphire coverslips and incubated for 5 to 10 min at 20°C prior to cryopreservation. As can immediately be appreciated on the representative electron micrographs, day-4 CTLs contain almost exclusively SCGs, while day-8 CTLs contained both MCGs and SCGs (Fig. 2A-B). The quantitative analysis reveals that the number of SCGs per cell and their density per cytoplasm area remained constant, whereas those of MCGs more than doubled between 4 and 8 days (Fig. 2C-D). The number of SMAPs per MCG did not vary during CTL expansion (Fig. 2E), confirming the measurement that each exocytosis event released a stable amount of SMAPs (Fig. 1G). Intriguingly, the SCGs, MCGs and SMAPs significantly grew in size over time in culture.

**Figure 2:**
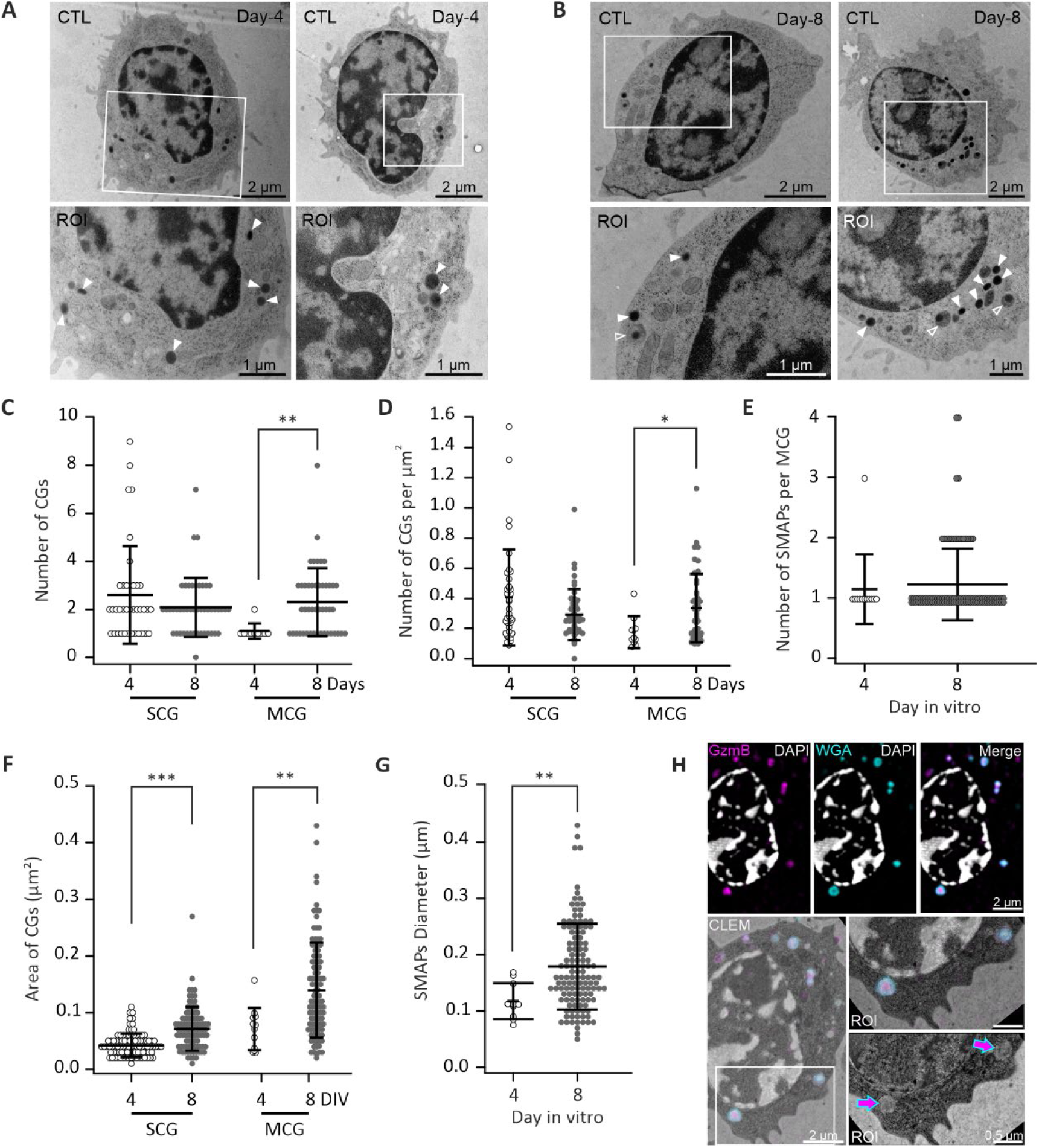
Post-activation time in vitro promotes multi-core granule (MCG) biogenesis in CTLs A-B. Display two representative day-4 (**A**) and 8 (**B**) CTLs. Regions of interest delineated by a white square on the top row are shown enlarged below. White filled arrowheads indicate single core granules (SCGs) and empty arrowheads point to MCGs. Scale bars are 2 µm on the top rows and 1 µm on the bottom row. **C-G.** Scatter dots plots showing the quantitative analysis of the transmission electron microscopy images of day 4 and 8 WT CTLs. The graphs display the total number of CGs per cell (**C**), the number of CGs per CTL cytoplasmic area (µm^2^) (**D**), and the number of SMAPs per MCGs (**E**). **F.** Shows the SCGs and MCGs area (µm^2^) and **G.** the SMAP diameter. Note that, due to space constraints, markers for one SMAP are staked on two rows in ***E***. All scatter dot plots show the mean and the SD as error bars. Statistical analyses were performed using Mann-Whitney Rank Sum Test. N_mouse_= 2, n_cells_ =27. Statistical significance is indicated as follows: \**p* < 0.05, \*\**p* < 0.01 and \*\*\**p* < 0.001. **H.** Representative correlative light and electron microscopy analysis of a day-6 GzmB-tdTomato (magenta) KI mouse CTL incubated with wheat germ agglutinin (WGA)-Alexa488 (cyan) to label the MCGs. Cells were seeded on anti-CD3ε antibodies coated coverslip for 10 min. prior to cryofixation. Shown are the super-resolved SIM images (top row) and the overlay with the corresponding electron microscopy image (bottom panel, left). Scale bar is 2 µm. In the magnified CLEM and TEM images (bottom panel, right), the magenta/blue arrows point to MCG intermediates that resemble multivesicular bodies, which contain GzmB. Scale bar is 1 µm.

Although MCG numbers increased significantly during in vitro expansion, we hypothesized that this alone could not account for the observed rise in SMAP-release events during culture. We therefore investigated MCG biogenesis by focusing on intermediate structures—organelles containing GzmB but lacking SMAPs and morphologically resembling multivesicular bodies (MVBs)— which might confound our quantification. Supporting this idea, CLEM performed by Chang, Schirra ^13^ had previously suggested the existence of such intermediates. To this end, we performed CLEM analysis of CTLs expanded for 6 days derived from GzmB–tdTomato knock-in mice. These cells secrete relatively low levels of SMAPs, consistent with incomplete MCG maturation. CTLs expressed fluorescently tagged GzmB and were co-labeled with wheat germ agglutinin conjugated to Alexa Fluor 488 (WGA–Alexa488), which selectively accumulates in biosynthetically active MCGs but not in SCGs (Cassioli et al., 2025; Chang et al., 2022). Using this approach, we confirmed the presence of MCG intermediates that do not contain fully formed SMAPs in day 6 CTLs (Fig. 2H).

### Immature multi-core granules are able to fuse at the immunological synapse

The presence of immature MCG intermediates raised the question of whether these organelles can fuse at the IS and, if so, whether they release their content as diffuse clouds or as discrete SMAPs. To address these questions, we used CTLs transfected with GzmB-pHuji and subjected to WGA-Alexa488 staining for 90 min prior to the experiment. CTLs were then seeded onto activating SLBs, and exocytosis of CGs was monitored by dual-color TIRF-microscopy for 15 min at 10 Hz and 20°C. We observed three distinct modes of secretion (Fig. 3A). The first corresponded to a cloud-release events that were not positive for WGA-Alexa488 (Fig 3A, top, Video S3). Given the specificity of WGA for MCGs, these events were attributed to the fusion of SCGs (Cassioli et al., 2025; Chang et al., 2022). The second corresponded to SMAP-release events, which as expected were labeled by WGA-Alexa488 (Fig 3A, middle, Video S4) and thus represent fusion of mature MCGs. In the third mode, we observed cloud-release events that were clearly associated with WGA-Alexa488 staining (Fig 3A, bottom, Video S5). We hypothesized that these events arise from the fusion of immature MCG structures, which we observed by CLEM (Fig. 2H). We first verified that the WGA staining did not change the overall behavior of the cells. We found that, as in unstained cells, the percentage of SMAP-release events increased significantly from 4 to 10 days (Fig. 3B), which was due to a concomitant reduction of the number of cloud-release events and increase in SMAP-release events (Fig. 3 C, D), with the rise in SMAP-release events being predominant. In addition, the number of SMAPs released per event was not changed over the time in culture (Fig 3. G). We next quantified the proportion of cloud-release events that were positive for WGA-Alexa 488. Approximately 40% of the cloud-release events were WGA-Alexa488 positive at day 4, decreasing to only about 10% at day 10 (Fig. 3E). This was not due to an increase in the number of WGA-Alexa488 negative events but rather a selective decrease in WGA-Alexa488 positive events (Fig. 3F). Collectively, these data indicate that the contribution of immature MCGs to fusion at the IS declines over time in culture as they mature into SMAP-containing MCGs.

**Figure 3.**
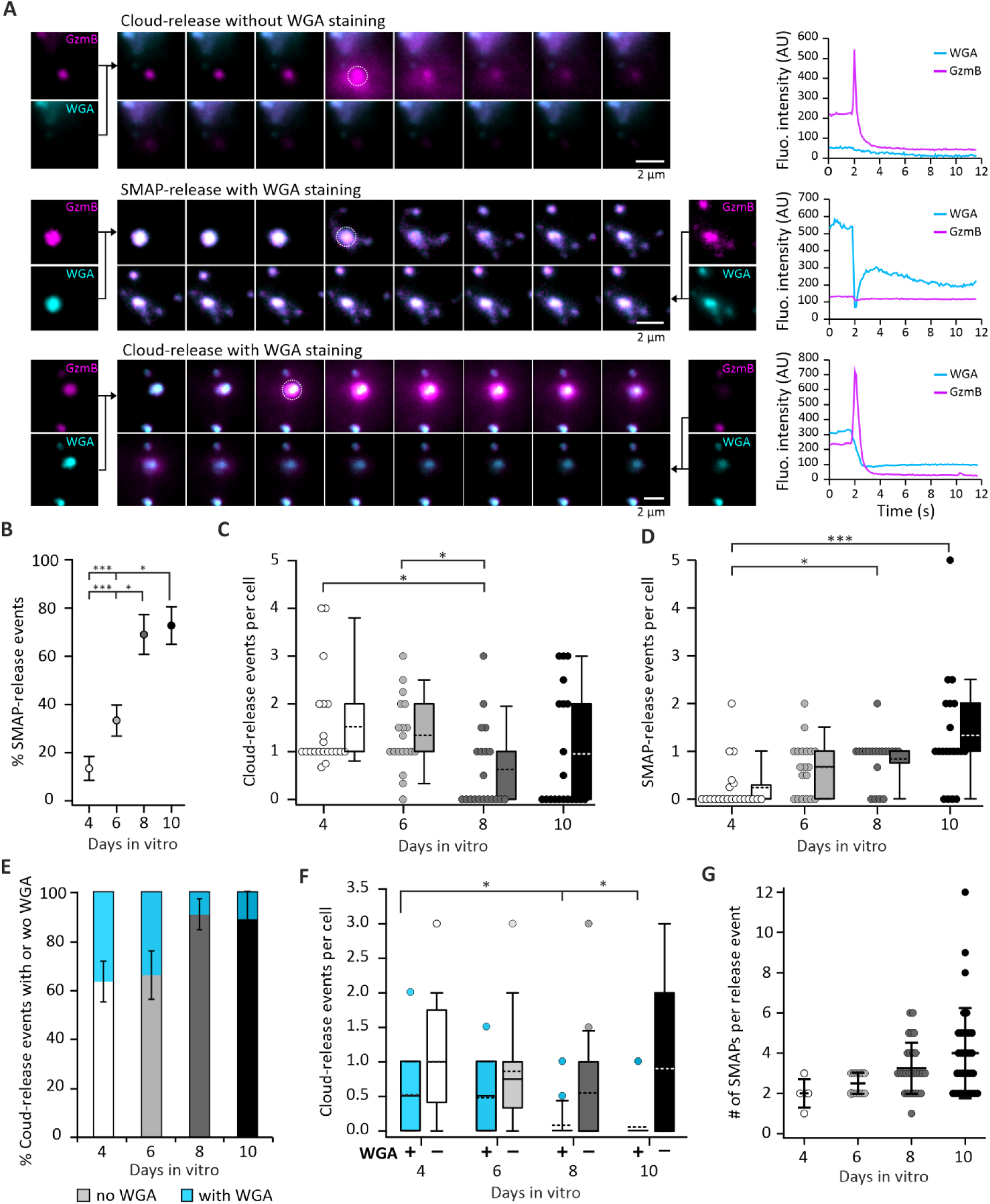
Cloud-release events can be associated with WGA labeling and decrease during in vitro expansion. **A.** Example of three types of exocytosis events recorded by TIRF-microscopy. SNAP shots of the event are displayed on the left and the fluorescence intensity over time of these events on the right. CTL overexpressing GzmB-pHuji (magenta) were stained with WGA-Alexa 488 (cyan), and seeded onto activating SLBs. Top row: Cloud-release event negative for WGA. GzmB release is fast and complete; Middle row: SMAP-release event positive for WGA. GzmB-pHuji and WGA signal decays slowly over time. The transient negative fluorescent spike corresponds to particle movement. Bottom: Cloud-release event negative for WGA. GzmB release is fast and complete, while WGA signal remains possibly due to staining of glycoproteins on exosomes. **B.** Diagram displaying the mean percentage of SMAP-release per CTL over their time in culture. Error bars are SEMs. **C-D.** Box and scatter dot plot of the normalized number of cloud- (**C**) and SMAP-release (**D**) per CTL and video over their time in culture. Stippled line in the box plot represents the mean and the solid line the median. **E.** Stacked bar graph representing the percentage of the cloud-release events that are WGA^+^ (cyan) or WGA^-^ (gray) over days in vitro. Error bars are SEM. **F.** Box plot of the normalized number of WGA^+^ (cyan) or WGA^-^ (gray) cloud-release) over days in vitro. The outliers are shown as symbols. **G.** Scatter dot plot showing the number of SMAPs released per exocytosis event over CTL time in culture. Shown are the mean and the SD as error bars. N_mice_ =5-6; n_video_= 21, 19, 20 and 20; n_cell_ = 31, 36, 25, and 27 for 4, 6, 8, and 10 days post-activation, respectively. Significance was tested with Kruskal-Wallis one-way ANOVA on rank followed by Dunn’s multiple comparison. * *p*<0.05, *** *p*<0.001.

We hypothesized that the GzmB present in immature MCGs slowly condensates in SMAPs. If that were the case, then we should detect this evolution through longer fluorescence decay times of the GzmB-pHuji signal in WGA-Alexa488 positive cloud-release events in comparison to the WGA-Alexa488 negative events. As can be appreciated by the fluorescent plot over time there was no obvious difference between cloud-release events with or without WGA-Alexa488 label (Fig. 3A right). In-depth quantitative analysis confirmed this observation: the average decay times were 2.88 ± 0.18 s and 3.10 ± 0.34 s, for WGA-Alexa488 negative and positive cloud-release events, respectively. In comparison, the average decay time of SMAP release events was 152.4 ± 21.6 s for SMAP-release events. Therefore, GzmB is probably in a similar diffusible state in SCGs and in immature MCGs. Intriguingly, the WGA staining remained high at the secretion site of immature MCG. This persistent signal is most likely attributable to WGA-labeled glycoproteins associated with exosomes present within MCGs ^6^.

### Thrombospondin-1 relocalizes to multi-core granules as they mature over time

TSP-1 contributes to the glycoprotein shell of SMAPs and integrates MCGs at a late phase of their maturation ^14^. This should be reflected in an increased colocalization of TSP-1 and MCGs over time. To assess this, CTLs isolated from GzmB–tdTomato mice and overexpressing TSP-1–GFPspark were co-stained with WGA–Alexa488 for 90 min, seeded onto poly-L-ornithine for 15 min, fixed with PFA, and imaged by structured illumination microscopy (SIM). Visual inspection of the images revealed a pronounced temporal shift in TSP-1 localization: at 4 days post-activation, TSP-1 displayed a mix of diffuse and punctate distribution poorly correlated with GzmB and WGA, whereas at 8 days post-activation it adopted a highly punctate pattern overlapping with GzmB- and WGA-positive structures corresponding to MCGs (Fig. 4A). To evaluate this phenomenon, we measured the coefficient of variation of the TSP-1-GFPSpark fluorescence. We found that at day 4 the coefficient of variation value was very low at 0.61 ± 0.02 denoting a diffuse signal, whereas at day 8 the value more than doubled to 1.36 ± 0.11 (N_mice_=4, n_cells_=32, 33, Fig. 4B), consistent with increased TSP-1 compartmentalization.

**Figure 4:**
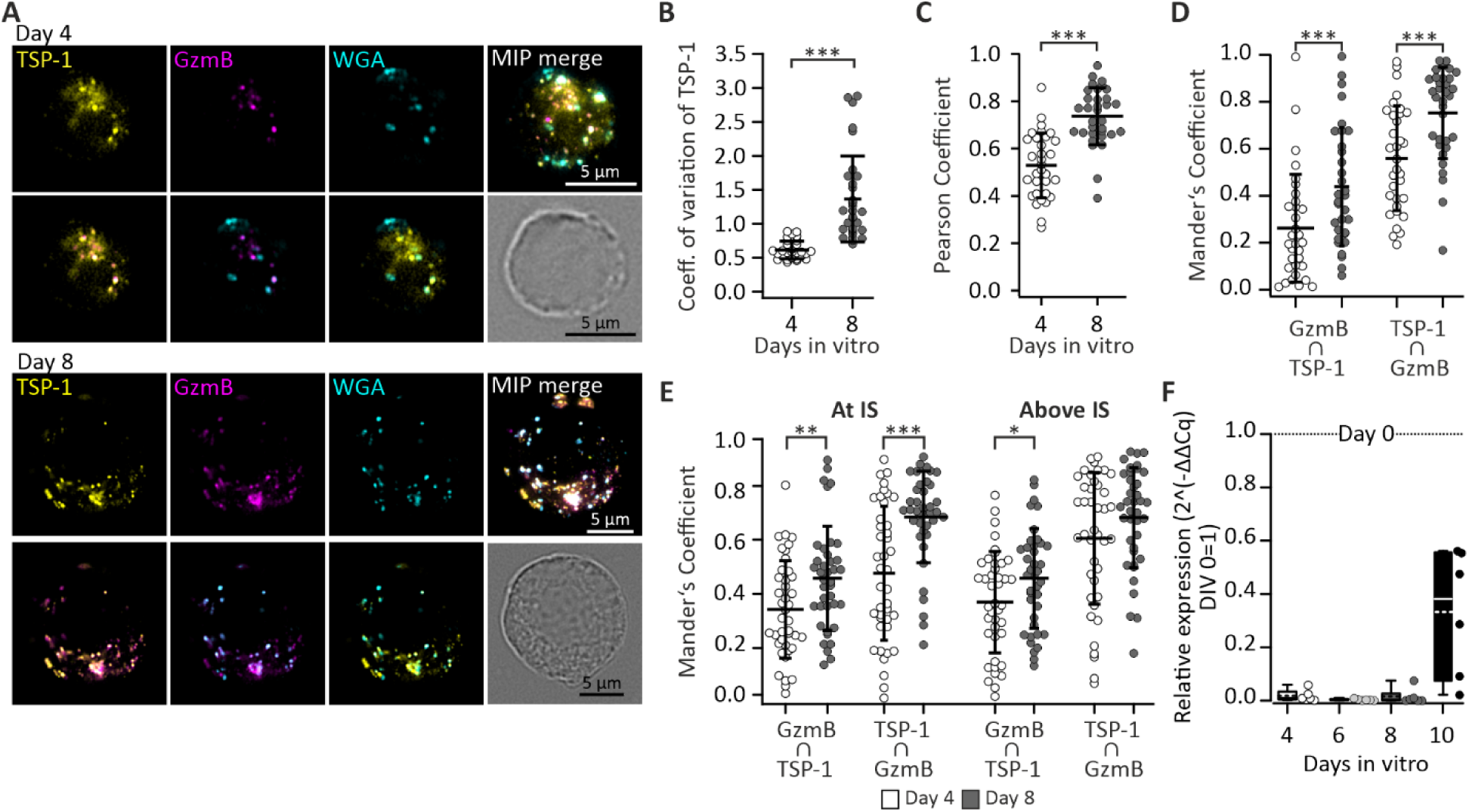
Dynamic redistribution of TSP-1 into multi-core granules over time in culture. **A.** Exemplary SIM images showing day-4 (top) and 8 (bottom) CTLs seeded on poly-L-ornithine coated coverslips, labeled with the endogenous GzmB-tdTomato fluorescence (magenta) and the overexpressed TSP-1-GFPSpark (yellow) and WGA-Alexa647 (cyan). Displayed are single plane images of the individual channels, (top row) and merged images of two channels at a time (bottom row). Finally, the maximum intensity projection of all merged channels is shown on the right with the bright field image of the cell below. Scale bar is 5 µm. **B.** The scatter dot plot shows the coefficient of variation (CV) of the TSP-1 signal from day 4 and 8 CTLs. CV = SD pixel fluorescence/mean fluorescence. **C.** Scatter dot plot representing the co-localization analysis of TSP-1 and GzmB with the Pearson’s correlation coefficient. **D.** Scatter dot plot displaying the degree of the overlap between GzmB-tdTomato and TSP-1-GFPSpark signals quantified with the Mander’s coefficient. **E.** Scatter dot plot showing Mander’s coefficient for GzmB-tdTomato and TSP-1-GFPSpark signals in CTLs stimulated with activating supported lipid bilayers (SLBs). The coefficients were measured within a zone of 1 µm of the immunological synapse (At IS) and in the rest of the cell (Above IS). Day-4 and 8 CTLs are shown in white and grey, respectively. The mean and the error bars representing SDs are shown in all scatter dot plots. Statistical analyses were performed using the Mann-Whitney Rank Sum Test. N_mouse_= 4, n_cells_=32 and 33 in the resting group seeded onto poly-L-ornithine coated coverslips, and n_cells_=39 and 41 in the stimulated group seeded on activating SLBs for day 4 and 8 CTLs, respectively. Statistical significance is indicated as follows: \**p* < 0.05, \*\**p* < 0.01 and \*\*\**p* < 0.001. **F.** Time course analysis of *THSP-1* mRNA levels in day 4 to 10 CTLs normalized to naïve T cells (day-0). The box and scatter dot plots show the relative abundance normalized to day 0 (control value = 1) of *THSP-1* transcripts. N_mice_ = 6.

We next quantified TSP-1 colocalization with GzmB using Pearson’s and Manders’ coefficients. Both analyses demonstrated a significant increase in TSP-1–GzmB colocalization with time in culture (Fig. 4C, D). To determine whether this redistribution was affected by cell stimulation, CTLs were seeded for 15 min in the presence of 10 mM calcium on anti-CD3ε monoclonal antibody–coated coverslips. Manders’ colocalization analysis again revealed a higher degree of TSP-1–GzmB overlap at day 8 compared to day 4. However, no significant difference was observed between regions within 1 µm of the IS and the remainder of the cell (Fig. 4E), indicating that increased colocalization occurs globally rather than being restricted to the synapse.

Finally, we assessed endogenous *THSP-1* expression dynamics by RT–qPCR. Consistent with previous observations in human CTLs ^14^, *THSP-1* transcript levels were strongly downregulated upon activation, decreasing by more than 10 times between naïve CD8⁺ T cells (day-0) and activated CTLs. However, *THSP-1* expression increased again by 10 days post-activation. This result suggests a conserved, maturation-dependent regulation of TSP-1 transcription.

The re-localization of TSP-1 from a diffuse signal to a punctate signal at MCGs is consistent with the hypothesis that SMAPs are generated late during the maturation of MCGs. Previous studies have shown that TSP-4 incorporation in MCGs precedes and is a prerequisite for TSP-1 to enter MCGs in order to generate SMAPs ^14^. Therefore, we sought to assess the contribution of TSP-4 to the maturation of MCG.

### Thrombospondin-4 over-expression promotes SMAP- over cloud-release events

We hypothesized that TSP-4 might be a critical factor that allows immature MCG to evolve in mature SMAP-containing MCG. To assess TSP-4 expression dynamics during CTL in vitro expansion, we analyzed *THBS-4* mRNA levels by RT–qPCR in CTLs 4, 6, 8, and 10 days post-activation (Fig. 5A). Similar to human CTLs, *THBS-4* expression was detectable at low levels in WT mouse CTLs at 4 days after activation and progressively increased over time in culture, reaching twice the level at day 10 compared to day 4.

**Figure 5.**
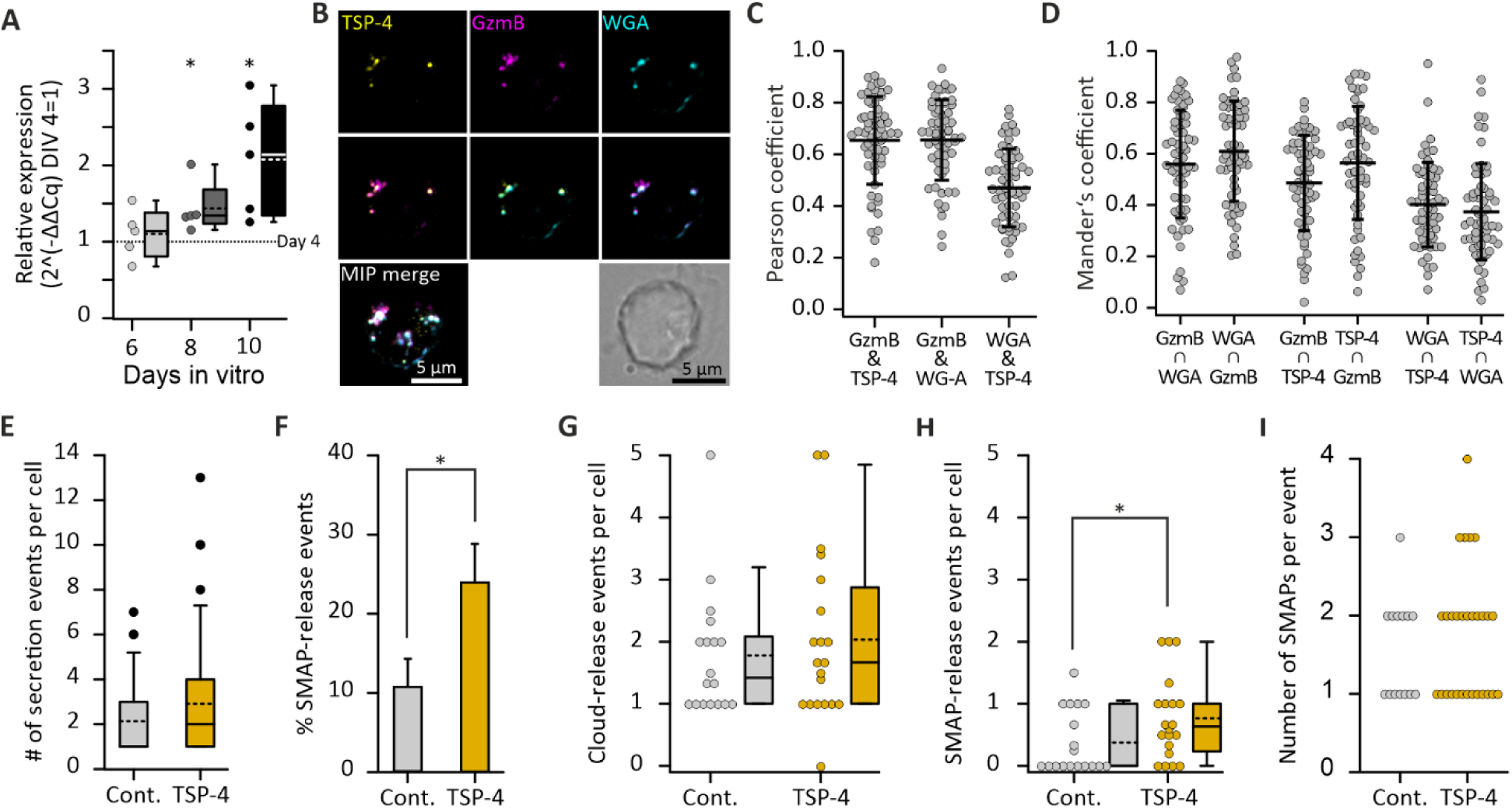
Overexpressed Thrombospondin-4 accumulates in multi core granules promoting SMAP-release. **A.** Time course analysis of *THBS-4* mRNA levels in day 6, 8 and 10 CTLs. The box and scatter dot plots show the relative abundance normalized to day 4 CTLs (ctr value = 1) of *THBS-4* transcripts. Statistical analysis was performed using the One-sample t-test and indicated as *p<0.05. N_mice_ = 5. **B.** Exemplary SIM images showing day-6 CTLs from GzmB-tdTomato KI mice (magenta), transfected with TSP-4-GFPSpark (yellow) and fed with WGA-Alexa647 (cyan) before seeding them onto anti-CD3ε antibody coated coverslips. Shown are single plane images of the individual channels (top row), and the merged images of two channels at a time (middle row). Finally, the maximum intensity projection image of all merged channels is shown on the bottom left with the bright field image of the cell on the right. Scale bar is 5 µm. **C.** Scatter dot plot showing the co-localization analysis using Pearson’s correlation coefficient. **D.** Scatter dot plot displaying the degree of the overlap between all three labels measured with the Mander’s coefficient. The mean and the error bars representing SDs are shown in all scatter dot plots. N_mouse_= 2, n_cells_=60. **E-I.** GzmB-tdTomato KI CTLs, transfected with GFP (control, white) or TSP-4-GFPSpark (TSP-4, green), were seeded onto activating SLBs and recorded with TIRF-microscopy. **E.** Box plot showing the number of secretion events per cell. The symbols show the outliers. **F.** Diagram displaying the mean percentage of SMAP-release per CTL. Error bars are SEMs. **G-H.** Scatter and box dot plot of the normalized number of cloud- (**G**) and SMAP-release (**H**) per CTL and video. The stippled line in the box plot represents the mean and the solid line the median. **I.** Scatter dot plot showing the number of SMAPs released per exocytosis event. N_mice_ =8; n_video_= 18 and 20; n_cell_ = 37 and 46 for control and TSP-4 overexpressing cells, respectively. Significance was tested with Mann-Whitney rank sum test. * *p*<0.05.

Next, we verified that over-expressed TSP-4-GFPSpark integrates MCGs at an early stage during their maturation. Day-6 CTLs isolated from GzmB-tdTomato KI mouse and transfected with TSP-4-GFPSpark were stained with WGA-Alexa647, then seeded on poly-L-ornithine coated coverslips for 15 min prior to PFA fixation and imaged by SIM. Visual inspection of the images revealed a punctate staining with high degree of colocalization between TSP–4 and the other two markers (Fig. 5B). Quantitative analysis confirmed the strong colocalization of TSP–4 with GzmB, as indicated by high Pearson’s (0.65 ± 0.02) and Manders’ (0.56 ± 0.03; N_mice_ = 2, n_cells_ = 60) coefficients, which exceeded the colocalization of TSP–4 and WGA (Fig. 5C, D). These data indicate that overexpressed TSP–4 associates with MCGs at an early stage of their maturation.

Given this early integration of TSP–4 into MCGs, we next asked whether TSP–4 overexpression would promote SMAP secretion in day 6 CTLs. CTLs from GzmB–tdTomato knock-in mice were transfected with constructs encoding either TSP–4–GFPSpark or GFP alone used as a control, then they were seeded on activating SLBs, and recorded by TIRF-microscopy. While TSP-4 over-expression did not affect the overall GzmB secretion nor the number of SMAP released per event (Fig. 5I), it significantly promoted the proportion of SMAP-release events (Fig. 5E, F). This effect was primarily driven by a significant increase in the number of SMAP-release events (Fig. 5G, H). These results indicate that TSP-4 accelerates MCG maturation.

### Cytotoxic T lymphocyte reactivation drives elevated SMAP release possibly through increased interferon-γ production

It was proposed that SMAPs are especially required to eliminate tumors in situations of transient and multiple interactions, where a delayed delivery of Gzms and Perforin might be advantageous to kill resistant cancer cells ^2, 12^. If that were the case, then it would be advantageous if several encounters with target cells would hasten MCG maturation thereby enhancing SMAP production and release. We investigated this possibility by restimulating day-5 GzmB-tdTomato-KI CTLs with fresh anti-CD3ε/anti-CD28 antibody coated beads for 4 h at 37°C. After a washing step the cells were kept in culture for another 12 h before using them in the experiment at 6 days post-activation. Control cells were not subjected to bead restimulation and also used at day 6. Secretion of CTLs seeded on activating SLBs was measured by TIRF-microscopy. We found that the restimulation protocol strongly enhanced the proportion of SMAP-release events (Fig. 6A). This shift was due to a concomitant significant decrease of cloud-release events and significant increase of SMAP-release events (Fig. 6B, C). Additionally, we observed that bead restimulation was the only treatment that significantly enhanced the number of secreted SMAPs per exocytosis event (Fig. 6D).

**Figure 6.**
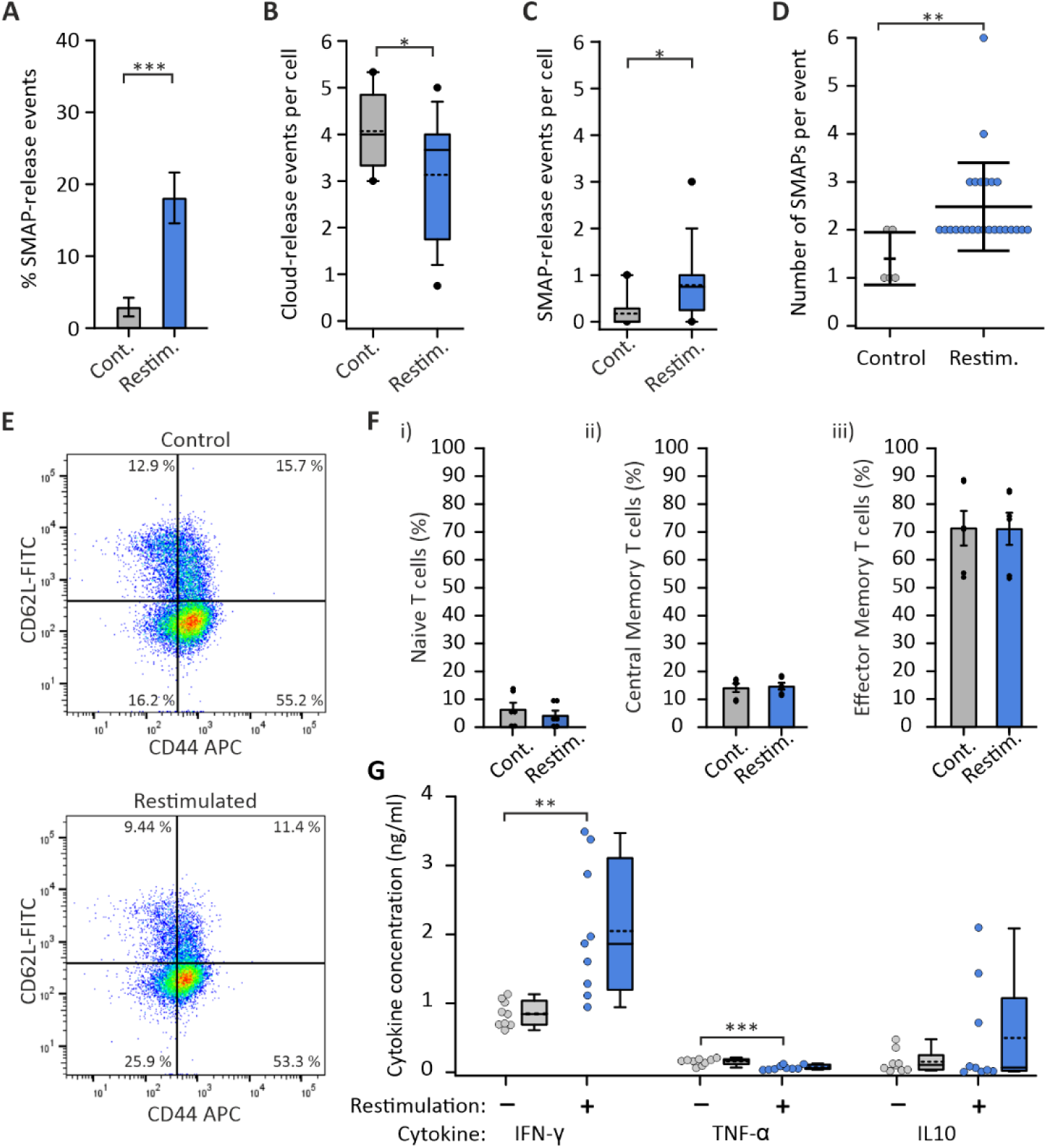
Cytotoxic T lymphocyte restimulation enhances secretion of SMAP and interferon-γ. GzmB secretion, cell differentiation state and cytokine release were compared between unstimulated control CTLs (gray) and CTLs restimulated on day 5 for 4 h with anti-CD3ε/anti-CD28 antibody coated beads (blue). Cells were measured about 12 h after restimulation on day 6. CTLs were isolated from GzmB-tdTomato KI mice. **A-D.** Cytotoxic granule exocytosis was monitored in CTLs seeded on activating SLBs using TIRF-microscopy over 15 min at 10 Hz. N_mice_ =4; n_video_= 9 and 15; n_cell_ = 34 for control and restimulated cells, respectively. **A.** Diagram displaying the mean percentage of SMAP-release per CTL. Error bars are SEMs. **B-C.** Scatter and box dot plot of the normalized number of cloud- (**B**) and SMAP-release (**C**) per CTL and video. Stippled line in the box plot represents the mean and the solid line the median. **D.** Scatter dot plot showing the number of SMAPs released per exocytosis event. Represented are the mean and the SD error bars. **E.** Representative dot plot showing the percentage of naïve (CD44-/CD62L+), central memory (CD44+/CD62L+), and effector memory (CD44+/CD62L-) CD8⁺ T cells in control (top) and restimulated condition (bottom). **F.** Statistical analyses of the restimulation effect on the CTLs subsets. The average percentages of (i) naïve T cells, (ii) central memory, and (iii) effector memory T cells are shown. Error bars are SEM. Individual values are displayed as scatter dot plot on top of the bars. N_mice_= 3. **G.** Scatter and box plot representing the concentration of IFN-γ, TNF-α and IL-10 quantified in culture supernatants derived from control and restimulated CTLs using a LEGENDplex™ MU Th1 panel. N_mice_=3, n_technical repeats_=9. Significance was tested with Mann-Whitney rank sum test. * *p*<0.05, ** *p*<0.01, *** *p*<0.001. Stippled line in the box plot represents the mean and the solid line the median.

We verified that this restimulation did not affect the differentiation of the CTLs by a flow cytometric analysis of the surface markers CD44 and CD62L (Fig. 6E), carried out on cells from identical cultures to those used for the TIRF-microscopy experiment. Overall, the proportion of naïve, central memory and effector memory CTLs was identical independently whether the cells were subjected or not to the 4 h-bead restimulation (Fig. 6F).

Stimulating CTLs with anti-CD3ε/anti-CD28 coated beads induces plethoric effects among which is the secretion of cytokines ^34, 35^, which may in turn regulate MCG maturation. To assess this, cytokine concentrations were measured by flow cytometry using the LEGENDplex™ MU Th1 (5-plex) panel in supernatants collected at day 6 from the same cultures used for the TIRF microscopy experiments. This assay enables the detection of a defined set of cytokines, including IL-2, IL-6, IL-10, TNF-α, and IFN-γ. Among these, only IFN-γ levels were increased in supernatants from 4 h-bead restimulated CTLs relative to unstimulated controls (Fig. 6G), suggesting a potential role for IFN-γ in MCG maturation. IL-6 concentration was below the detection limit under both conditions. A decreasing trend in TNF-α was observed upon 4 h-bead restimulation, but TNF-α concentrations was below the quantification limit of 0.1 ng/ml (Fig. 6G). No significant differences were detected in IL-10 levels between the two conditions (Fig. 6G). Quantification of IL-2 was confounded by the presence of recombinant IL-2 in the culture medium used for CTL expansion.

### Interferon-γ promotes multi-core granule maturation via upregulation of *THBS-4* expression

Our data shows that CTL restimulation enhances release of both SMAPs and IFN-γ. To determine whether these events are independent or if IFN-γ promotes MCG maturation and thereby augments SMAP secretion, we neutralized IFN-γ in the culture medium using 5 µg/ml anti-IFN-γ antibody starting 2 days post-activation until the day of analysis. We then compared GzmB release with TIRF-microscopy at day 4 and 8 in treated and untreated CTLs seeded onto activating SLBs. While IFN-γ neutralization had no effect at 4 days post-activation, the increased proportion of SMAP-release events observed at day 8 in control CTLs was abolished upon treatment (Fig. 7A). This effect was not attributable to changes in the overall number of cloud-release events, but rather to a significant reduction in SMAP-release events at 8 days post-activation in cells treated with neutralizing anti-IFN-γ antibodies compared to controls (Fig. 7B, C). Finally, *THBS-4* mRNA levels were significantly reduced in day 6 and 8 CTLs cultured in the presence of neutralizing anti-IFN-γ antibodies compared to control CTLs (Fig. 7D), as assessed by RT-qPCR. All together, these results indicate that IFN–γ promotes MCG maturation and SMAP secretion, at least in part, through regulation of *THBS–4* expression.

**Figure 7.**
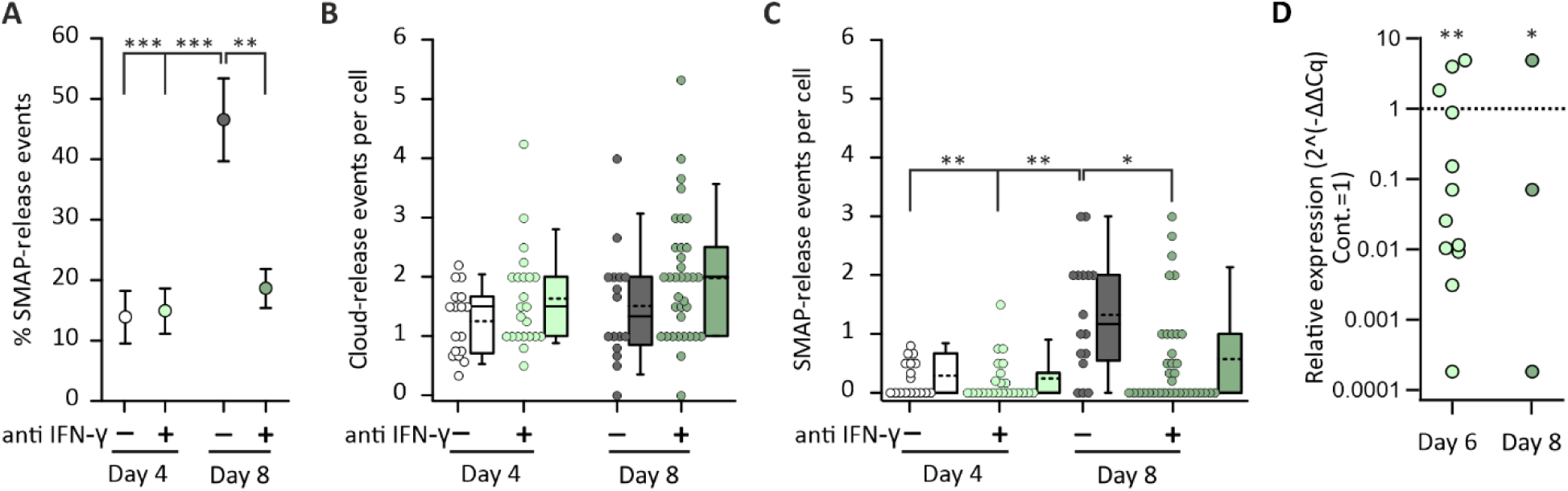
Interferon-γ controls MCG maturation through *THBS-4* expression. WT CTLs were treated with (+, green) or without (-, gray, control) 5 µg/ml anti-IFN-γ antibody starting 2 days post-activation throughout the rest of the culture time by supplementing the medium each day with the antibody during cell splitting. **A-C.** CTLs transfected with GzmB-pHuji were seeded onto activating SLBs and imaged for 15 min at 10 Hz by TIRF-microscopy. **A.** Diagram displaying the mean percentage of SMAP-release per CTL. Error bars are SEMs. **B-C.** Scatter and box dot plot of the normalized number of cloud- (**B**) and SMAP-release (**C**) per CTL and video. Stippled line in the box plot represents the mean and the solid line the median. N_mice_ =2 - 4; n_video_= 17 and 23; n_cell_ = 45 and 63 for day 4 control and treated cells, respectively and n_video_= 16 and 35, n_cell_ = 34 and 79 for day 8 control and treated cells, respectively. Significance was tested with Mann-Whitney rank sum test. * *p*<0.05, ** *p*<0.01, *** *p*<0.001. **D.** Analysis of *THBS-4* mRNA levels in day 6 and 8 CTLs cultured in the absence (control) or presence of neutralizing anti-IFN-γ antibodies. The box and scatter dot plots show the relative abundance of *THBS-4* transcripts in treated CTLs normalized against control CTLs (control value = 1). * *p*<0.05, ** *p*<0.01, Student t-test was performed using ΔCq values. N_mice_ = 12 and 3 for day 4 and 8, respectively.

## Discussion

Murine and human CTLs contain different CGs, in which the lytic substances Perforin and Gzms are found either in a diffusive form in SCGs or in a stabilized form as SMAPs in MCGs ^12, 13, 14^. In the present work we addressed the question whether the secretion of SCGs vs. MCGs can be differentially regulated. Using TIRF-microscopy, that allowed us to observe in real time the exocytosis of both CG types, we found that in vitro CTL expansion was the main factor to enhance SMAP-release (Fig. 1, Fig. 8A) and that it could be recapitulated by short term restimulation (Fig. 6). To understand the underlying mechanism, we performed a TEM analysis of young (4 days post-activation) and old (8 days post-activation) CTLs and observed that while the number of SCGs remained constant the number of SMAP containing MCGs increased significantly over time (Fig. 2). An additional CLEM analysis of CTLs revealed the existence of MCG intermediates (Fig. 2H). These were characterized as organelles containing intraluminal vesicles, together with the cytotoxic effector GzmB and the MCG specific marker WGA, but lacking SMAPs. We further demonstrated that these MCG intermediates labelled with WGA were capable of exocytosis and that they released GzmB in a diffusive manner (Fig. 3, Fig. 8A). Their number diminished over time in culture in favor of fully mature MCGs that liberated GzmB as SMAPs. This transition of MCG intermediates to fully mature MCG was promoted by IFN-γ (Fig. 7, Fig. 8A) possibly via *THBS-4* gene expression upregulation (Fig. 5, Fig 7D). Finally, using a calcein based killing assay we could show that CTLs, which were maintained in culture for long time, killed significantly better than younger cells (Fig. 1 H, I). The difference in killing capacity was only observed after long contact time (> 7 h) between CTL and target cells and at high E:T ratio (10:1) suggesting that the difference was due to the enhanced SMAP release in older CTLs.

**Figure 8.**
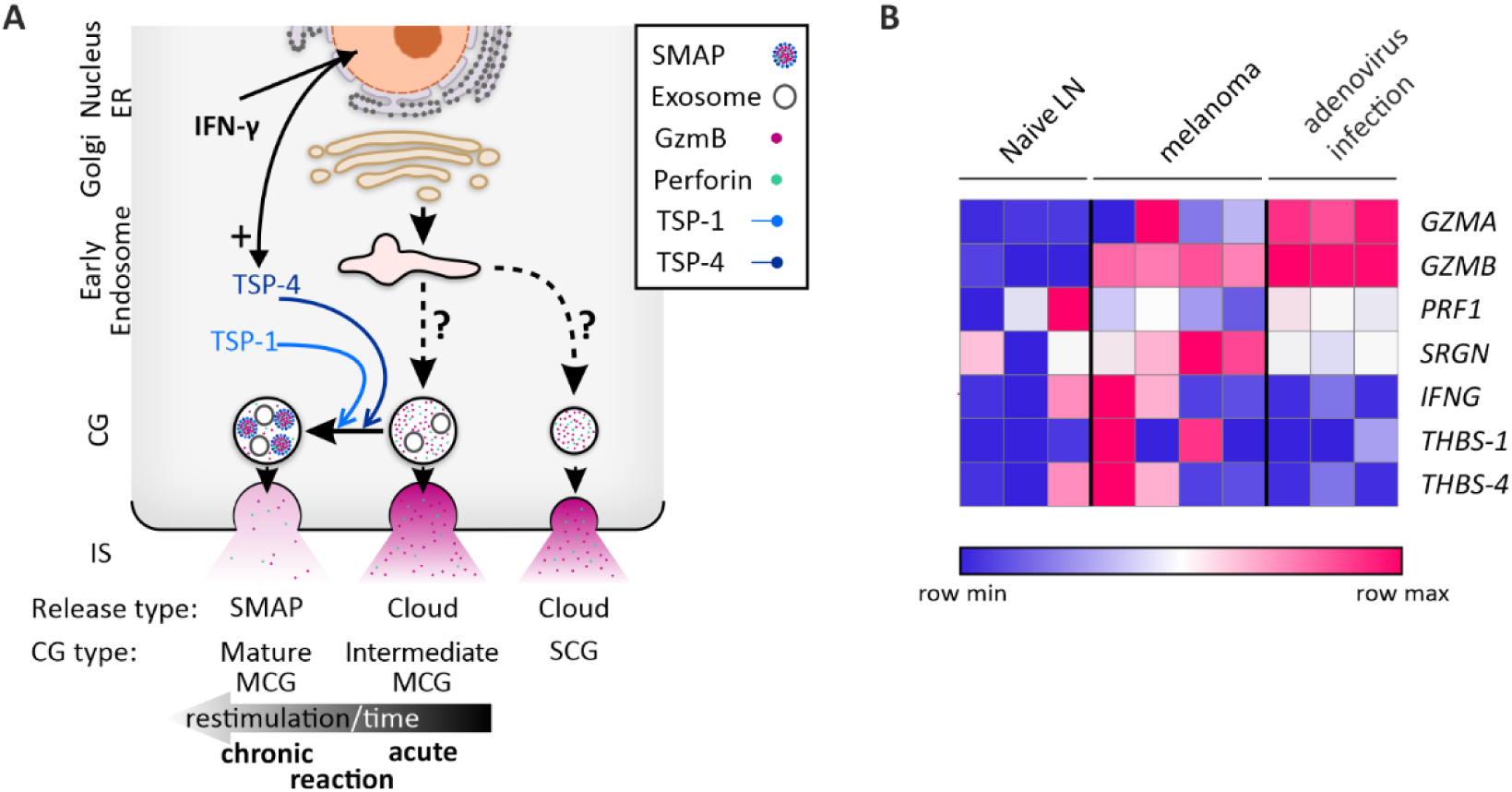
IFN-γ–dependent maturation of cytotoxic granules drives SMAP formation and defines context-specific CTL effector program. **A.** Shortly after activation CTL produce SCGs and intermediate MCGs that release their content as diffusive material. If the target cells are eliminated quickly, such as in acute virus infections, then intermediate MCGs do not reach fully mature state. Upon longer activation time, such as in a chronic infection or cancer, IFN-γ level rises, inducing enhanced TSP-4 transcription and MCG maturation. TSP-4 enters MCG followed by TSP-1 allowing the generation of SMAPs which are released upon target cell contact. **B.** Transcriptomic profiling of cytolytic machinery and SMAP structural components in murine CD8^+^ T cells under different conditions. Row-normalized expression heatmap of primary murine CD8^+^ T cells isolated from naive lymph nodes (n=3), autochthonous melanoma lesions (n=4), and acute adenovirus infected lung tissue (n=3), mined from repository dataset E-GEOD-42824. While classical cytolytic effectors (Granzymes – *GZMA*, *GZMB*; Perforin – *PRF1*; Serglycin – *SRGN*) are globally upregulated upon both viral and malignant challenge, the Supramolecular Attack Particle (SMAP) core structural scaffolds Thrombospondin-1 (*THBS-1*) and Thrombospondin-4 (*THBS-4*) display a distinct, heterogeneous enrichment pattern specific to a subset of the exhausted tumor-infiltrating lymphocyte (TIL) microenvironment. Baseline variance of *THBS-4* is noted within a single naive control replicate. The color bar denotes relative expression levels from row minimum (blue) to row maximum (red).

The observed shift in secretion type over time suggests that cell differentiation could contribute to this process. Between 4 and 10 days post-activation, the proportion of CTLs secreting SMAPs increased from 8.2% to 64.6% (Fig. 1). Over the same period, the proportion of effector memory cells increased by only 1.6-fold (Fig. S2). Similarly, restimulation of CTLs with anti-CD3ε/anti-CD28–coated beads increased the fraction of SMAP-secreting cells from 14.7% in control cultures to 52.9%, while the distribution of CTL subtypes remained unchanged (Fig. 6). These observations indicate that the transition of MCGs from an intermediate to a mature, SMAP-bearing form is independent of cell differentiation.

We showed that CTLs contain SCGs, MCGs and MCG intermediates and that all three types of granules can fuse at the IS formed on activating surfaces. Such heterogeneity is consistent with previous studies demonstrating that CGs can assume multiple morphologies ^36, 37, 38, 39^, including intermediate forms detected by CLEM analysis ^13^. Beyond morphological heterogeneity, it has been shown that CG subtypes also differ in their molecular composition and release dynamics. In this context, GzmB has been demonstrated to colocalized with CD63, whereby CD63 is localized to the organelle membrane and the intralumenal vesicle they may contain ^6, 40^. By studying the fluorescence decay rate of CD63-pHuji and GzmB-superecliptic pHluorin, Alawar, Schirra ^6^ found that CD63 could be associated with either SCGs (fast fluorescence decay) or with MCGs (slow fluorescence decay). At 5 days post-activation, ∼75% of exocytotic events were attributed to MCG fusion; however, a large proportion of these MCGs exhibited rapid GzmB decay (Fig. 2G of Alawar, Schirra ^6^) indicative of MCG intermediates.

The question that arises is how the MCG intermediates can be exocytosis competent. The fusion competence of the CG is given by the presence of a v-SNARE on its membrane, which is VAMP2 for murine CTLs, its association with Munc13-4, Munc18-2 and the presence of t-SNAREs, SNAP23 and Syntaxin11 on the IS membrane. The accumulation of the t-SNAREs in the plasma membrane is induced by the IS formation and is independent of the CG ^41^. A colocalization study of VAMP-2 with GzmB using a VAMP-2mRFP knock-in mouse showed that nearly all GzmB positive granule also contained VAMP2 ^13, 42^. It is thus likely that MCG intermediates also bear VAMP2 on their surface. Due to the transient nature of priming/docking factors localization on CGs at the time of exocytosis it is more challenging to verify the association of MCG intermediates with Munc13-4 and Munc18-2. However, a very high degree of colocalization of Munc13-4 and GzmB at the IS has been shown ^43^. Therefore, all exocytosis components appear to be reunited to allow MCG intermediates to be secreted.

Our data revealed that TSP-4 overexpression accelerates MCG maturation and that *THBS-4* expression increases during cell expansion. Consistently, neutralization of IFN–γ reduces both MCG maturation and TSP–4 expression. Previous studies have shown that IFN–γ induces *THBS-4* expression in macrophages, promoting a pro–inflammatory phenotype ^44^. Together, these findings suggest that IFN–γ promotes the transition of MCG intermediates into mature MCGs by inducing *THBS-4* expression. The molecular mechanisms linking IFN–γ signaling to *THBS-4* transcription are not well understood. IFN–γ activates multiple transcriptional pathways, including STAT1 via GAS elements, as well as the mTOR/PI3K/AKT and MAPK pathways that converge on transcription factors such as CREB and MYC ^45^. However, while little is currently known about the regulation of the *THBS-4* promoter, changes in proteins expression in inflammation and tissue injury due to post-transcriptional mechanisms have been reported ^46, 47^. In silico analyses identify CREB, among others, as a potential transcriptional regulator of *THBS-4* (https://www.genecards.org), which is an important regulator of CD8 T cell differentiation ^48^.

The real–time calcein release–based killing assay revealed that CTLs maintained in culture for 8 days post-activation kill B16 melanoma cells more efficiently than CTLs at day 4. SMAPs are highly stable and retain cytotoxic activity for hours to days following their release ^12^. It has therefore been proposed that SMAP–mediated killing proceeds on a longer timescale than cytotoxicity mediated by diffusible Perforin and Gzms, and that SMAPs may bypass synaptic resistance mechanisms ^2, 3, 12^. We leveraged these properties to distinguish SMAP–dependent killing from other cytotoxic mechanisms. First, we employed B16V melanoma cells, which display limited sensitivity to Fas/FasL–mediated apoptosis due to reduced Fas expression ^49, 50, 51^, thereby biasing cytotoxicity toward Perforin– and Gzm–dependent pathways. Furthermore, melanoma cells have been reported to be relatively resistant to soluble Perforin and Gzms ^32, 33^, suggesting that they may be preferentially susceptible to SMAP–mediated killing. In addition, we extended the assay duration to 7 hours, rather than the commonly used 4 hours, to allow sufficient time for SMAP–dependent cytotoxic effects to manifest. Up to 3.5 hours, the killing efficiency of day 4 and 8 CTLs was indistinguishable. Beyond this time point, the killing curves gradually diverged, with the enhanced cytotoxic capacity of day 8 CTLs becoming significant at 7 hours at an E:T ratio of 10:1. A similar trend was observed at an E:T ratio of 1:1; however, in this case the difference between day 4 and day 8 CTLs did not reach statistical significance. Recent evidence indicates that exosomes contained within CGs can exert potent cytotoxic activity ^6^. However, because these exosomes are already present in both intermediate and mature MCGs (this work and ^6^) and are therefore expected to be released at comparable levels by day 4 and day 8 CTLs, they are unlikely to account for the late divergence in killing efficiency observed here. Taken together, our findings strongly support the conclusion that the enhanced killing capacity of CTLs 8 days after activation relative to 4 days is attributable to increased SMAP secretion.

An important unresolved question is the functional relevance of the slow, time-dependent transition of MCGs toward the formation of mature SMAPs. Acute viral infections are typically controlled within days, and CTLs engaged in this context are likely activated only transiently. Moreover, virus-infected cells generally lack robust immune evasion mechanisms and may therefore be efficiently eliminated without the need for SMAP-mediated cytotoxicity. Conversely, chronic viral infections, some intracellular bacterial infections that are causative of tuberculosis, and tumor development occurs over weeks to years. Chronic microbial infections diverge rapidly from acute infections due to co-evolved mechanisms ^52^. On the other hand, cancer cells frequently acquire immune evasion strategies that confer resistance to diffusible Perforin and Gzms ^53^, but potentially not to SMAPs. CTLs operating in the tumor microenvironment are exposed to prolonged antigenic stimulation and repetitive restimulation, conditions that as we demonstrated favor enhanced SMAP biogenesis. These observations suggest that time- and stimulation-dependent SMAP formation may represent an adaptive mechanism that enables CTLs to tailor their cytotoxic arsenal to the nature and persistence of the pathological challenge. This hypothesis is supported by our meta-analysis of repository transcriptomic data (Fig. 8B). While acute adenovirus infection induces robust upregulation of classical cytolytic effectors and IFN-γ, it completely fails to induce the SMAP structural scaffolds *THBS-1* and *THBS-4*. In contrast, CTLs isolated from autochthonous melanoma lesions display a distinct, albeit heterogeneous, enrichment of both *THBS-1* and *THBS-4* transcripts (Fig. 8B). Importantly, a detailed understanding of the kinetics of SMAP formation will be critical for the rational design of SMAP-based or SMAP-enhancing immunotherapeutic strategies.

## Materials and Methods

### Mice

All the experimental procedures were conducted in compliance with the regulations of the state of Saarland (Landesamt für Verbraucherschutz, AZ.: 2.4.1.1 and 11/2021). Wild-type (WT) mice were purchased from Charles River, and GzmB-tdTomato knock in (KI) mice ^54^ were purchased from the Transgenesis and Archiving of Animal Models (TAAM) (National Centre of Scientific Research (CNRS), Orleans, France). Transgenic mice and WT mice used in this study were all in C57BL/6N background. Additionally, all the animals used for experiments were at the age of 8-13 weeks-old and of both sexes. They were kept under specific pathogen free (SPF) conditions at 22 °C room temperature with 50–60% humidity and 12 h dark/light cycles. They were provided with food and water ad libitum.

Mice were anesthetized with CO_2_ and executed by cervical dislocation. Splenocytes were isolated as previously described ^55^. Briefly, the left abdominal cavity was exposed, and the spleen was carefully removed, placed on a 70 μm cell strainer (Corning Life Sciences, Singapore), and ground. The grinding slurry remaining on the strainer was rinsed with RPMI medium, and the cell suspension (10 ml) was collected into a 15 ml sterile centrifuge tube (Corning Life Sciences) and centrifuged (6 min, 1100 rpm). Washed splenocytes were mixed and incubated with 1 ml erythrocyte lysis buffer (155 mM NH_4_Cl, 10 mM KHCO_3_ and 0.13 mM EDTA, pH 7.3) for 30 s ^56^. To terminate lysis, 10 ml of ice-cold RPMI was added and the cells were twice centrifuged (6 min, 1100 rpm). Primary CD8^+^ T lymphocytes were positively isolated from spleens according to the instructions of the Dynabeads FlowComp Mouse CD8 Kit (Thermo Scientific, Waltham, MA, USA).

### Cell culture

The isolated naive CD8^+^ T cells were stimulated with anti-CD3ε/anti-CD28 activator beads (1:0.8 ratio) and cultured for up to 9 days at 37 °C with 5% CO2. Cells were cultured at a density of ∼1 × 10^6^ cells/ml in a 24-well plate (Greiner Bio-One) with AIM-V medium (Invitrogen) supplemented with 10% FCS, 1% penicillin‒streptomycin (Invitrogen). On day 2 poststimulation, 100 U/mL recombinant IL-2 (Gibco) was added to support T-cell proliferation. The resulting activated effector CTLs were used for subsequent experiments.

B16-V murine melanoma cells (DSMZ-Leibniz Institute, Germany) were cultured in RPMI 1640 (Gibco, Thermo Fisher Scientific) supplemented with 10% FBS under standard conditions (37 °C, 5% CO₂). Cells were passaged every 2–3 days at a 1:10 ratio. For passaging, cells were detached using trypsin/EDTA (∼5 min), collected, centrifuged, and reseeded in fresh complete medium.

### Electroporation of CTLs

Mouse CTLs were electroporated at the indicated time points using a Mouse T Cell Nucleofector Kit (Lonza) with 2.5 µg plasmid DNA (pMAX-GzmB-L-pHuji, pMAX-TSP-1-GFPSpark, or pMAX-TSP-4-GFPSpark) using the Amaxa Nucleofector system (program X-001) as described by Alawar, Schirra ^57^. Following electroporation, cells were transferred into pre-warmed recovery medium (OptiMEM-based transfection medium supplemented with 10% FCS, 10 mM HEPES, 1% DMSO, and 1 mM sodium pyruvate) and incubated at 32 °C for 14–18 hours. Cells were subsequently washed and cultured in IL-2–supplemented medium at 37 °C until use (∼2 hours). Electroporated CTLs were used for imaging of granule secretion at the immunological synapse by TIRF-microscopy.

### Treatment of CD8^+^ T cell culture with neutralizing anti-IFN-γ antibodies

Isolated CD8⁺ T cells were cultured in AIM V medium supplemented with 10% FCS, 50 µM β-mercaptoethanol, and 100 U/mL IL-2. On day 2, cultures were divided into control and treatment groups. The treatment group was supplemented daily with anti-mouse IFN-γ antibody (Ultra-LEAF™, BioLegend, 1:200), whereas control cells were maintained under identical conditions without antibody. Cells from both groups were electroporated with GzmB-pHuji on days 3 and 7 and analyzed for exocytosis by TIRF-microscopy on days 4 and 8.

### CTL restimulation

Day 5 bead-activated GzmB-tdTomato KI CTLs were harvested, and the original activation beads were removed using a magnet. Cells were counted and restimulated with fresh anti-CD3ε/CD28-coated beads at a ratio of 1:0.8 in AIM V medium supplemented with 10% FCS and 50 µM β-mercaptoethanol (BME), in the absence of IL-2, for 4 h at 37 °C. Following restimulation, beads were removed, and cells were washed twice with warm imaging buffer by centrifugation at 900 rpm for 6 minutes. Cells were subsequently resuspended in fresh AIM V medium supplemented with 10% FCS, 50 µM BME, and 100 U/mL recombinant mouse IL-2, and incubated for an additional 12 hours at 37 °C and 5% CO₂. Cells were then used for downstream applications, including TIRF-microscopy or flow cytometry.

### Activating supported lipid bilayers (SLB)

SLBs were prepared for TIRF-microscopy imaging to visualize granule secretion at the immunological synapse. SLBs were prepared as previously described ^58, 59^. Briefly, the glass coverslips that were washed with acid piranha and subjected to plasma cleaner, were mounted on sticky-Slide VI0.4 (Ibidi) to form 6 flow channels. Small unilamellar liposomes were prepared using 18:1 DGS-NTA(Ni) (790404C-AVL, Avanti Polar Lipids), 18:1 Biotinyl Cap (870282C-AVL, Avanti Polar Lipids), and 18:1 (D9-Cis) PC (850375C-AVL, Avanti Polar Lipids) in specific mixtures at total lipid concentration of 4 mM. SLBs were formed by incubating 50 µL of the liposome suspension in each flow channel for 20 min at 20 ± 2 °C. SLB were then washed with HEPES buffer containing 1 mM CaCl_2_ and 2 mM MgCl_2_ (HBS) and blocked with 1% human serum albumin (HBS/HSA) prior to functionalization. This occurred in two steps. Initially, the SLBs were treated with 5 µg of streptavidin per channel (S11226, Thermo Fisher Scientific) for a period of 10 min. at a temperature of 20±2°C. This was followed by three washings. Finally, 30 µg per channel of biotinylated monoclonal anti-mouse CD3ε antibody (BD Pharmingen, clone 145-2C11) was linked to the streptavidin on SLB, and 50 µg/ml of 12-histidine-tagged mouse ICAM-1 was linked to nickel ions.

### Total internal reflection fluorescence (TIRF) microscopy

Secretion was measured for CTLs that were maintained in culture for 4, 6, 8, or 10 days depending on the experimental setup. They were either isolated from WT mice and transfected with GzmB-L-pHuji alone or additionally stained with WGA-Alexa488 (1 µg/mL, 1.5 h at 37 °C, Lonza). Or else they were isolated from GzmB-tdTomato mice and used as such or transfected with TSP-4-GFPSpark.

For imaging, cells were allowed to settle for 1–2 minutes on activating SLBs (BD Pharmingen, clone 145-2C11) at 10 µg/ml, unless otherwise stated. The cells were then perfused with extracellular buffer containing calcium (10 mM glucose, 5 mM HEPES, 140 mM NaCl, 4.5 mM KCl, 2 mM MgCl₂, and 10 mM CaCl₂) to stimulate CG secretion. Cells were recorded for 15 min. at 20 ± 2 °C at a frequency of 10 Hz with a 100 ms exposure time. The TIRF-microscopy setup (Visitron Systems GmbH) was based on an IX83 microscope (Olympus) equipped with a UAPON100XOTIRF NA 1.49 objective (Olympus), solid-state excitation lasers at 488 nm, 561 nm, an iLAS2 illumination control system (Roper Scientific SAS), an Evolve-EM 515 camera (Photometrics), and a filter cube containing Semrock FF444/520/590/Di01 dichroic and FF01-465/537/623 emission filters. The system was controlled via Visiview software (version 4.0.0.11; Visitron Systems GmbH).

Time-lapse series were analyzed via ImageJ (version 1.54p) or the FIJI package for ImageJ and the IVEA software tool ^60^. For cells transfected with GzmB-pHuji a fusion event was defined by a sharp fluorescence intensity increase followed by a diffusion cloud alone (cloud-release) or by remaining fluorescent particles (SMAP-release). For cells derived from GzmB-tdTomato mice, the exocytosis event was accompanied by smaller raise in fluorescent intensity as the probe is not pH sensitive ^60^.

Fusion was always determined using the GzmB channel. If a co-labelling was performed then the other channel was only used for colocalization study. Fusion parameters were quantified as follows: the average percentage of SMAP-release was determined relative to the total number of fusion events per cell; the number of cloud- or SMAP-release events per cell correspond to the number of events per video normalized to the number of secreting cell (i.e., each dot in these scatter dot plots correspond to one experimental repeat).

### Killing assay

Cytotoxic activity was evaluated using a calcein release assay. B16V target cells were labeled with 500 nM Calcein-AM in serum free AIM-V medium supplemented with 10 mM HEPES and incubated for 15 min at room temperature protected from light. Cells were washed then seeded at 2.5 × 10⁴ cells per well in black 96-well plates with clear bottoms (Greiner BIO-ONE, Cat # 655892). After seeding, target cells were allowed to settle in the plate for 15–20 min at 37°C prior to the addition of effector cells. CD8⁺ T cell effector populations were prepared by removal of CD3ε/CD28 stimulation beads, followed by centrifugation and resuspension in AIM-V/HEPES medium. Effector and target cells were co-cultured at effector-to-target (E:T) ratios of 1:1 and 10:1 in triplicate wells. Appropriate controls included target cells cultured alone to determine spontaneous release, target cells treated with 1% Triton X-100 (final concentration) to determine maximal lysis, and medium-only wells for background fluorescence correction. Anti-mouse CD3ε antibody was added to effector cells immediately prior to co-culture initiation. Fluorescence measurements at the bottom of the wells were acquired every 10 min. for 8 h at 37°C using a GENios Pro microplate reader (Tecan, excitation: 485 nm; emission: 535 nm). Percent target cell lysis was calculated by normalizing experimental fluorescence values to spontaneous and maximal release controls following subtraction of background fluorescence using the following equation:

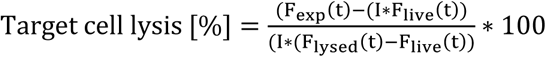

Whereby F_exp_(t) is the fluorescence of the tested sample at time t, F_lysed_(t) is the fluorescence of the lysed control at time t, F_live_(t) is the fluorescence of the live control at time t, and I is the Index at time 0 (F_exp_(0) / F_live_(0)).

### Flow cytometry analysis

For flow cytometry analysis was performed at days 4, 6, 8, and 10 depending on the experimental setup. 0.5 × 10⁶ CD8^+^ T cells without activator beads were resuspended in D-PBS (Gibco) and incubated in the dark on ice for 30 min. with the following cell surface markers: anti-CD25-PE (BD Pharmingen, clone PC61, 1:400), anti-CD62L-FITC (BD Pharmingen, clone MEL-14, 1:200), and anti-CD44-APC (BD Pharmingen, clone IM7, 1:200). Data acquisition was performed on a BD FACSAria III analyzer (BD Biosciences) via BD FACSDiva™ software. Flow cytometry data were analyzed with FlowJo v10.0.7 software.

### Multiplex cytokine analysis (LEGENDplex™)

Extracellular cytokines (IL-2, IL-6, IL-10, TNF-α, and IFN-γ) were quantified using a LEGENDplex™ Mouse Th1 Panel (5-plex) (BioLegend) according to the manufacturer’s protocol. Supernatants were diluted and incubated with fluorescent capture beads, followed by biotinylated detection antibodies and streptavidin-PE. Samples were analyzed on a BD FACSAria III flow cytometer (BD Biosciences), and cytokine concentrations were determined using standard curves with LEGENDplex™ Data Analysis Software (BioLegend). Cell supernatant was collected from the same cultures as those used for TIRF-microscopy experiment on day-6, with cells that have been subjected to the restimulation protocol or not (control).

### RNA isolation, cDNA synthesis, and RT–qPCR

WT CD8^+^ T lymphocytes (5 × 10^6^ cells) were collected from cell culture without the activator beads. Total RNA was extracted via TRIzol reagent (Thermo Fisher Scientific) following the manufacturer’s protocol. Briefly, 5 × 10⁶ CD8⁺ T cells were homogenized in 500 µl of TRIzol reagent, followed by the addition of 100 µl of chloroform. The samples were subsequently centrifuged at 11,400 × g for 15 minutes at 4 °C, after which the mixture was separated into three distinct phases, with the RNA remaining exclusively in the upper aqueous phase. The aqueous phase was transferred to a fresh tube and the RNA was precipitated by mixing with an equal volume of 100% isopropanol. The RNA pellet was washed once with 75% ethanol, air-dried, and dissolved in DEPC-treated H₂O. The quality and quantity of the extracted RNA were assessed via a NanoDrop spectrophotometer.

For *THBS-1* RT qPCR, the isolated RNA was subsequently reverse transcribed into complementary DNA (cDNA) via the SuperScript™ II RT (Thermo Scientific) in accordance with the manufacturer’s instructions. Briefly, 1 µg of RNA was mixed with a reaction mixture containing random primer mixture, 10 mM dNTPs, and the SuperScript™ II enzyme. The mixture was first incubated at 25 °C for 10 minutes, followed by incubation at 42 °C for 50 min. Enzyme inactivation was achieved by heating the mixture to 70°C for 15 minutes. RT‒qPCR was performed via the use of Genious 2X SYBR Green Fast qPCR Mix (ABclonal) on a Bio-Rad CFX96 Touch Real-Time PCR Detection System. A total of 10 ng of cDNA was used as a template in each reaction. Amplification was performed according to the manufacturer’s protocol via specific primers targeting. *THBS-1* (forward: 5′-TGTGAGGTTTGTCTTTGGAA-3′, reverse: 5′-ACCATGCTGGATAGTTCATC-3′) and IFN-γ (forward: 5′-GAACGCTACACACTGCATCT-3′, reverse: 5′-GTCACCATCCTTTTGCCAGT-3′). TATA-box binding protein (TBP) (Mm_Tbp_1_SG /QT00198443) was used as an internal reference gene to normalize target gene expression. Relative gene expression levels were quantified via the 2^(-ΔCt).

*THBS-4* RT-qPCR was performed to quantify relative THBS-4 transcript levels in CD8⁺ T cells across different days in vitro (Fig. 5A) and in the presence or absence of anti–IFN-γ antibodies (Fig. 7D). Total RNA was reverse-transcribed into complementary DNA (cDNA) using either the iScript cDNA Synthesis Kit (Bio-Rad) or the ABScript II First Strand cDNA Synthesis Kit (ABclonal), respectively. For each reaction, 1 µg of RNA was used per sample, and cDNA synthesis was performed according to the manufacturers’ instructions. For Fig. 5A, the mixture was incubated 5 min at 25 °C for primer annealing, followed by 20 min at 46 °C for cDNA synthesis and 1 min at 95 °C for enzyme inactivation. For Fig. 7D, the reaction mixture was incubated 5 min at 25 °C for primer annealing, followed by a 60 min incubation at 42 °C for cDNA synthesis, and 5 min at 80 °C for enzyme inactivation. Quantitative PCR for Fig. 5A was carried out using the Sso Fast™ Eva Green® Super Mix (Bio-Rad) and the CFX Duet Real-Time PCR System (Bio-Rad), under standard cycling conditions with optimized annealing temperatures for the primer pairs used. Each reaction contained 50 ng of cDNA template. Amplification of a cDNA fragment corresponding to *THBS-4* transcript was performed using the following primers: forward 5′-GCCCAGGAAGATAGCAACAA-3′ and reverse 5′-TGATGTGTGCCCTGAGAATG-3′. Quantitative PCR for Fig. 7D was carried out using the same qPCR Mix (ABclonal) and detection system as for *THBS-1* (described previously), with each reaction containing 25 ng of cDNA template. Relative *THBS-4* mRNA expression was determined on triplicate samples using the 2^(−ΔΔCq) method ^61, 62^ and normalized to *HPRT1* (hypoxanthine-guanine phosphoribosyltransferase) mRNA levels. Statistical Student’s t-test was performed with the ΔCq values. Raw data were analyzed using the CFX Maestro 2.3 software (Bio-Rad) (for Fig. 5A).

### Electron Microscopy and Post-embedding CLEM

For ultrastructural analysis day 4 and day 8 WT mouse CTLs were seeded on poly-L-ornithine and anti-CD3ε coated sapphire discs in flat specimen carriers for 10 min at 37°C and 5% CO2, enabling the formation of an immunological synapse. Cells were vitrified in a high-pressure freezing system (EM PACT2, Leica) in RPMI medium with 30% FCS and 10 mM HEPES. Freeze substitution and embedding in Epon 812 epoxy resin (Electron Microscopy Sciences) was performed as previously described ^63^. All samples were processed in an automatic freeze-substitution apparatus (AFS2, Leica). Briefly, samples were placed into the precooled (−130 °C) freeze-substitution chamber of the AFS2 and the temperature was linearly increased from −130 to −90 °C over 2 h. Cryo-substitution was performed in 2% osmium tetroxide in anhydrous acetone and 2% water. The temperature was increased linearly from −90 °C to −70 °C over 20 h, from −70 °C to −50 °C over 20 h, and from −50 °C to −10 °C over 5 h. After washing with anhydrous acetone, the cells were embedded in Epon-812 (30% Epon/acetone for 15 min at −10 °C, 70% Epon/acetone for 1 h at −10 °C and pure Epon for 1 h at 20 ± 2 °C; Electron Microscopy Sciences). The temperature was increased linearly from 20 ± 2 °C to 60 °C for 4 h, and Epon was polymerized at 60 °C for 24 h. Following polymerization, carriers and sapphire discs were removed from the resin block. Ultrathin (70 nm) sections were cut perpendicular to the former sapphire surface using an ultramicrotome (EM UC7; Leica), collected on Pioloform-coated copper grids, stained with uranyl acetate and lead citrate and analyzed with a Tecnai12 Biotwin electron microscope (Thermo Fisher Scientific). The transmission electron microscopy (TEM) images were acquired using Olympus iTEM 5.0 image software (build1243). Only cells with intact and well-preserved membranes were included in the analysis.

Secretory lysosomes (SCGs), multivesicular cytotoxic granules (MCGs), and SMAPs were identified based on ultrastructural features. SCGs were defined as small, homogeneous electron-dense granules, whereas MCGs exhibited heterogeneous, multivesicular morphology. Quantification was performed manually using ImageJ, with regions of interest defined for cellular compartments and granule populations. The number of SCGs and MCGs per cell, as well as SMAPs per MCG, was determined, and SMAP size was quantified using Feret’s diameter.

To localize proteins within CTL organelles, post-embedding CLEM analysis of cryo-fixed GzmB-KI mouse CTLs incubated with WGA 488 was conducted as previously described (Chang et al., 2022). Day 6 CTLs seeded on poly-L-ornithine and anti-CD3ε coated sapphire discs in flat specimen carriers for 10 min at 37°C and 5% CO2, enabling the formation of an immunological synapse. Cells were vitrified as described above. For cryo-substitution, samples were transferred into the pre-cooled (−130°C) chamber of the AFS2 (Leica), where the temperature was increased gradually from −130 to −90°C over 2 h. Cryo-substitution was conducted at −90°C to −70°C for 20 h in anhydrous acetone and at −70°C to−60°C for 24 h with 0.3% uranyl acetate in acetone. Samples were infiltrated with increasing concentrations (30%, 60%, and 100%) of Lowicryl K11M/HM20 mixture for 1 h each, followed by 5 h of 100% Lowicryl infiltration. Samples were then UV polymerized at −60°C for 24 h and at 5°C for an additional 15 h. After polymerization, the membrane carriers were removed, and ultrathin sections (100 nm) were cut using an EM UC7 ultramicrotome (Leica), mounted on carbon-coated 200-mesh copper grids (Plano), and stored at 4°C until SIM and TEM analysis. Fluorescence analysis of EM grids was performed within 24 h post-sectioning to prevent loss of fluorescence signals. Grids were stained with DAPI (1:1000) for 3 min, washed, and sealed with silicone (Picodent Twinsil®) for SIM imaging. After fluorescence imaging, grids were stained with uranyl acetate and lead citrate for electron microscopy, captured with the Tecnai12 Biotwin transmission electron microscope (Thermo Fisher Scientific). Only CTLs with intact membranes, organelles, and nuclei were analyzed for correlation. DAPI images (405 nm) were used to align fluorescence with electron microscope images, allowing precise localization of fluorescent signals within cells. Image overlays were performed in Corel DRAW X6.

### Structured illumination microscopy (SIM)

For correlative light and electron microscopy (CLEM) imaging, ultrathin sections of post-embedded GzmB-mTFP KI mouse CTLs stained with WGA-Alexa488 were imaged using 488 and 561 nm wavelengths. SIM imaging with a 63× Plan-Apochromat objective (NA 1.4) covered a 200-mesh grid (around 90 µm²), ensuring precise alignment with the grid bars. DAPI images were taken at 405 nm to identify CTL nuclei and define the image plane. Z-stacks were acquired with a step size of 100 nm, and image processing was performed using Zen 2011 (Zeiss).

For colocalization analysis, CTLs from days 4 and 8 of GzmB-tdTomato KI mice were transfected with TSP-1-GFPSpark or TSP-4-GFPSpark, stained with WGA-Alexa488 (1 µg/mL, 1.5 h at 37 °C, Lonza) and seeded onto poly-L-ornithine– or anti-CD3ε antibody–coated coverslips (monoclonal anti-mouse CD3ε antibody (BD Pharmingen, clone 145-2C11)). Cells were incubated in low Ca²⁺ medium followed by Ca²⁺ stimulation (10 min.) and fixed with 2% paraformaldehyde. Samples were mounted in Mowiol and imaged using an ELYRA PS.1 system (Zeiss) equipped with a 63× Plan-Apochromat objective (NA 1.4). Z-stacks were acquired with a step size of 200 nm using 488, 561 and 647 nm excitation wavelengths. Image reconstruction was performed using ZEN software (Zeiss). For colocalization analysis, the JACoP plugin ^64^ of Fiji ^65^ was used. Pearson’s and Manders’ overlap coefficients ^66^ were used for quantification of the degree of colocalization. The coefficient of variation (CV) of the TSP-1-GFPSpark signal was measured to quantify the degree of protein clustering ^67^.

### Statistical analysis

All statistical analyses were performed using Igor Pro (v6.37) or SigmaPlot (v14.0). Data are presented as mean ± SEM (or SD, where indicated). Statistical tests used for each experiment are specified in the corresponding figure legends. Comparisons between two groups were performed using Student’s t-test or the Mann–Whitney U test, as appropriate. For multiple-group comparisons, one-way ANOVA followed by Tukey or Student–Newman–Keuls post hoc tests, or Kruskal–Wallis tests followed by Dunn’s multiple comparison test were applied. All tests were two-tailed with a 95% confidence interval. One-sample t-test was used to compare the mean of multiple samples to a reference value of 1, assigned to the control sample for normalization. Statistical significance was denoted as *p < 0.05, **p < 0.01, and ***p < 0.001. All imaging data were analyzed with ImageJ or FIJI (version 1.54p). Colocalization analyses were carried out using the JaCoP plugin, and Pearson’s and Manders’ coefficients were calculated.

## Author Contributions

Methodology and Investigation O.M.K., M.S., C.S.; E.L., S.-M.T., and C.C.; Data curation and formal analysis O.M.K., M.S., C.S.; L.S., S.-M.T., C.C. and U.B.; Resources and supervision H.-F. C., C.T.B., and U.B.; Manuscript writing U.B.; Manuscript revision H-F.C., C.S., M.L.D., C.T.B., and U.B.; Conceptualization, project administration, and funding acquisition H.-F-C., M.L. D., C.T.B., U.B.

## Acknowledgements

We thank Elmar Krause for support in imaging techniques (SFB894 (P1)). We are grateful to S. Valitutti and M.L. Dustin for critical comments on our manuscript. We thank Margarete Klose, Anja Bergsträßer, Nicole Rothgerber, and Katrin Sandmeier for excellent technical assistance.

## Ethics Statement

All experimental procedures were approved and performed in accordance with the regulations of the state of Saarland (Landesamt für Verbraucherschutz, AZ: 2.4.1.1).

## Data Availability Statement

Data and information that support the findings of this study will be available on Zenodo upon acceptance with restricted access. The data will be made available by the corresponding author upon reasonable request.

