## Supplementary Figure 1 and Supplementary Figure 2 for "Interferon-γ promotes SMAP production by cytotoxic T lymphocytes in a thrombospondin-4 dependent manner"

### Supplementary data

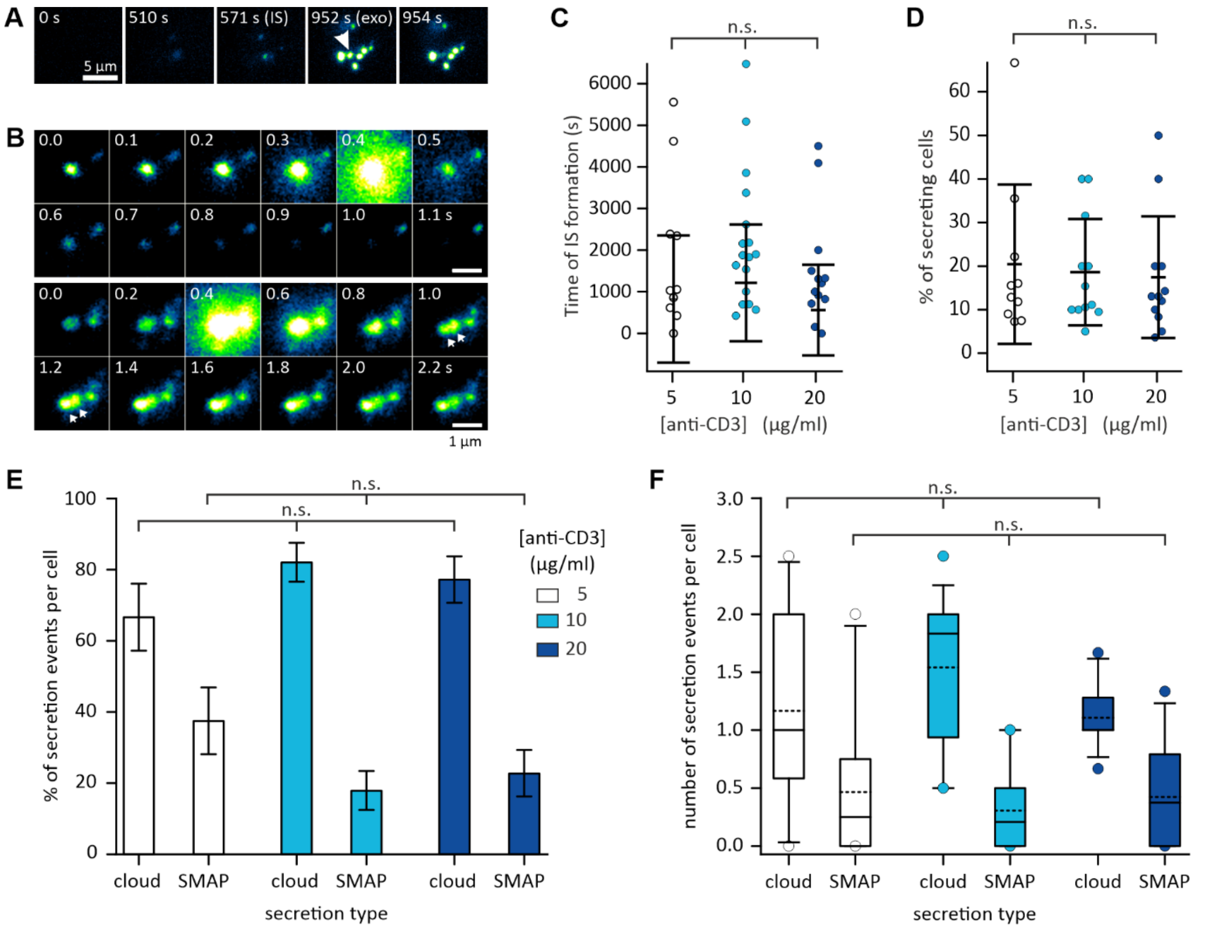


**Figure S1: Stimulation strength does not influence secretion type in cytotoxic T lymphocytes (CTLs)**
**A.** Snapshots of a CTL overexpressing GzmB-pHuji with total internal reflection fluorescence microscopy (TIRFM). Cytotoxic granule (CG) secretion was stimulated by seeding the cell on 10 µg/ml anti-CD3 containing supported lipid bilayer (SLB). Shown are the time of immunological synapse (IS) formation (571 s), the time just before (952 s) and after (954 s) CG fusion (arrow). Scale bar is 5 µm. **B.** Snapshots of various fusion types. Top row: cloud-release. Fusion of a CG, in which GzmB-pHuji is released as diffusible form characterized by a fluorescent cloud that quickly dissipate. Bottom row: SMAP-releases. Fusion of a CG, in which part of GzmB-pHuji remains aggregated as SMAPs (arrows) for several seconds after release. Time point of fusion is 0.4 s. Scale bars are 1 µm. **C-D.** Scatter dot plots of (**C**) the time of IS formation and (**D**) the percentage of secreting cells. Error bars are SDs. **E.** Bar diagram of the percentage of cloud- and SMAP-release per cell. CTLs were stimulated by SLBs containing 5 (white), 10 (cyan) and 20 (blue) µg/ml anti-CD3. Error bars are SEMs. **F.** Box plot of the normalized number of cloud- and SMAP-release per CTL and video depending on an the anti-CD3 concentration in the SLBs. $N_{mice}$ = 5, $n_{cell}$ = 23, 36 and 20 for 5, 10 and 20 µg/ml anti-CD3, respectively. n.s. corresponds to not significant.

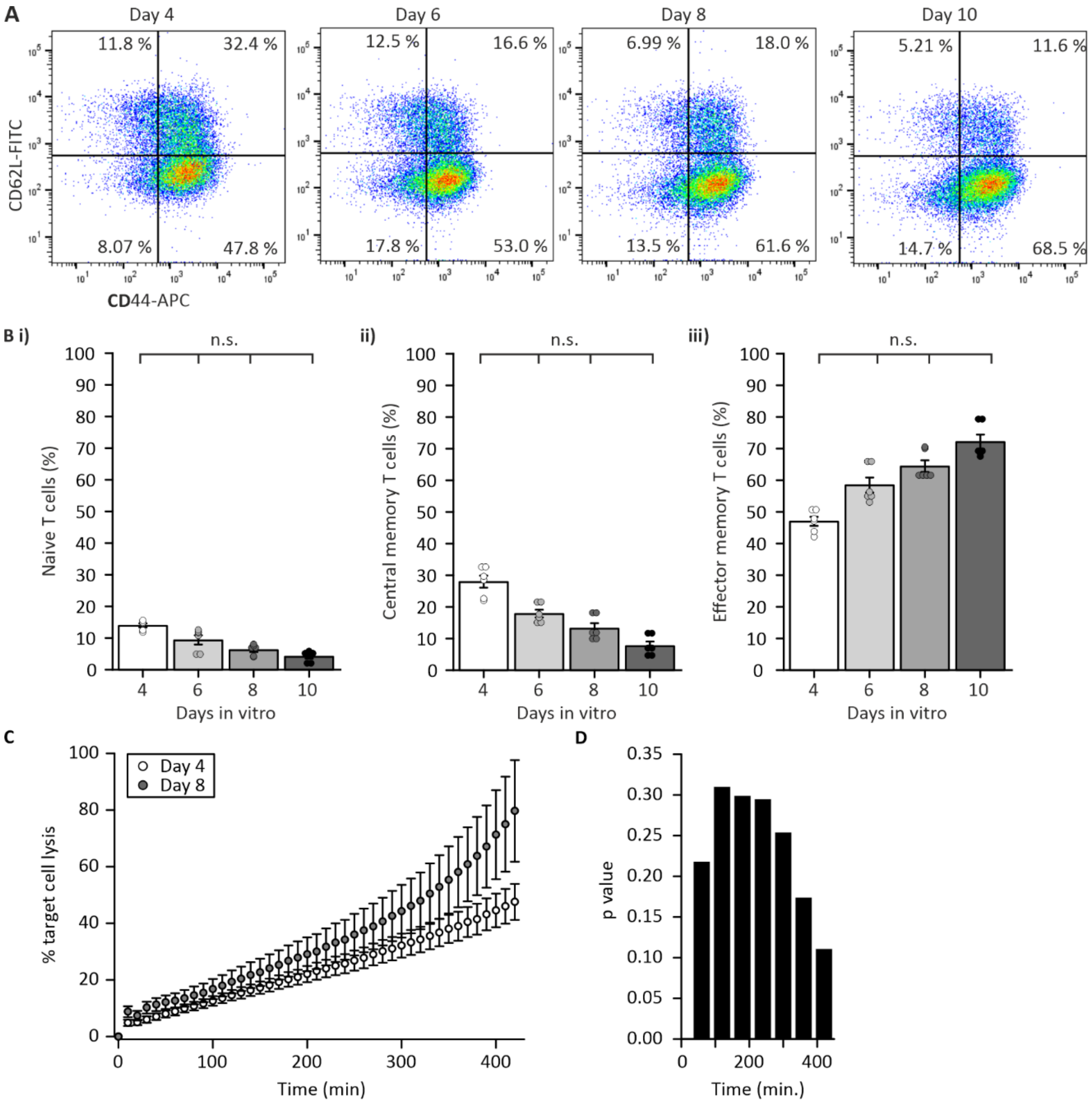


**Figure S2 (related to Figure 1): Changes of CTL subtypes does not explain increased SMAP-release over time in culture.**

**A.** Representative dot plot showing the percentage of naïve (CD44-/CD62L+), central memory (CD44+/CD62L+), and effector memory (CD44+/CD62L-) WT CD8$^+$ T cells on days in vitro 4, 6, 8, and 10. **B.** Statistical analyses of the effect of time in culture on the subtypes of CD8$^+$ T cells. Shown are the average percent of (i) naïve T cells, (ii) central memory, and (iii) effector memory T cells. Error bars are SEM. Individual values are shown as scatter dot plot on top of the bars. $N_{mouse}$= 3. These are a subset of the cultures shown in Figure 1. Statistical significance is indicated as follows: n.s. = not significant with Kruskal-Wallis test. **C.** Real-time calcein release-based killing assay. Day-4 (white) and day-8 (gray) CTLs were co-cultured with B16V cells at a 1:10 cell ratios, and killing was measured every 10 min for 7 h. Shown is the mean with error bars as SEMs. $N_{mice}$=3, $n_{repeats}$=9. **D.** Graph representing the p value of t-tests comparing the difference of killing percentage between DIV 4 and 8.

**Legend of the supplementary Videos.**

**Video S1** (related to Fig. 1A)**: Exemplary cloud-release events**
4 DIV old CTL expressing GzmB-pHuji displaying several cloud-release events. The last event is shown in the Fig. 1A, top row. Video was acquired at 10 Hz and is displayed at 200 Hz.

**Video S2** (related to Fig. 1A)**: Exemplary SMAP-release events**
10 DIV old CTL expressing GzmB-pHuji displaying several cloud-release events. The last event is shown in the Fig. 1A, bottom row. Video was acquired at 10 Hz and is displayed at 200 Hz.

**Video S3** (related to Fig. 3A)**: Exemplary cloud release events negative for WGA-Alexa488**
4 DIV old CTL expressing GzmB-pHuji (magenta), stained with WGA-Alexa488 (cyan) displaying several, indicated by arrows, cloud-release events that are negative for WGA. The last event is shown in the Fig. 3A, top row. Note that other events occur as well. Video was acquired at 10 Hz and is displayed at 200 Hz. Both channels (GzmB-phuji and WGA-Alexa488) were acquired simultaneously.

**Video S4** (related to Fig. 3A)**: Exemplary SMPA-release events positive for WGA-Alexa488**
10 DIV old CTL expressing GzmB-pHuji (magenta), stained with WGA-Alexa488 (cyan) displaying one SMAP-release events, indicated by an arrow, that is positive for WGA. The last event is shown in the Fig. 3A, middle row. Video was acquired at 10 Hz and is displayed at 200 Hz. Both channels (GzmB-phuji and WGA-Alexa488) were acquired simultaneously.

**Video S5** (related to Fig. 3A)**: Exemplary cloud-release events positive for WGA-Alexa488**
6 DIV old CTL expressing GzmB-pHuji (magenta), stained with WGA-Alexa488 (cyan) displaying one cloud-release events, indicated by an arrow, that is positive for WGA. The last event is shown in the Fig. 3A, bottom row. Video was acquired at 10 Hz and is displayed at 200 Hz. Both channels (GzmB-phuji and WGA-Alexa488) were acquired simultaneously.
